# Biotope-dependent High Level Resistance to Reactive Oxygen Species, Antibiotic Tolerance, and Virulence of *Staphylococcus aureus*

**DOI:** 10.1101/2025.10.31.685911

**Authors:** Vincent Léguillier, Karine Gloux, Majd Khalife, Rochelle D’Mello, Zoran Minic, Milica Sentic, Marta Catto, Marisa Manzano, Christine Péchoux, Sandrine Truchet, Philippe Gaudu, Christina Nielsen-Leroux, Brahim Heddi, Alexandra Gruss, Jasmina Vidic

## Abstract

The human and livestock pathogen *Staphylococcus aureus* poses a major clinical challenge due to antibiotic treatment failure. Its resilience is mainly attributed to antibiotic resistance and tolerance mechanisms related to persistence. Here we investigate how two infection-relevant biotopes, milk and serum, shape *S. aureus* pathogenic properties and capacity to withstand environmental stresses. Milk-*versus* serum-adapted bacteria show gross differences in envelope physical properties, membrane fatty acid composition and rigidity, and pigment production, and display distinct proteomic profiles. Compared to serum, milk adaptation of *S. aureus* confers extreme resistance to ROS damage, pronounced antimicrobial tolerance, and accelerated killing in an insect infection model. High level *S. aureus* pigmentation in whole milk is stimulated by milk lipids, and is responsible for high ROS resistance. The remarkable robustness of *S. aureus* in a milk biotope may signal the need to adjust antibiotic regimens when treating mastitis infections in humans and livestock.

## Introduction

The opportunist pathogen S*taphylococcus aureus* causes a wide array of life-threatening diseases that can affect multiple organs^1^. Methicillin-resistant *S. aureus* (MRSA) was first reported in 1961 in the United Kingdom in a clinical isolate and now accounts for over 60% of *S. aureus* isolates in US hospitals^2^. A decade later, the first foodborne MRSA strain was found in milk from Belgian cows with mastitis^3^. MRSA Staphylococci are now common pathogens isolated from mastitis milk and pose a significant health concern in dairy cattle^4,5^. The recovery time after infection can be long in cows because *S. aureus* can persist in teat canals, teat lesions, and mammary glands. During this period, milk is unsafe for human consumption and infected animals can serve as a reservoir for staphylococcal transmission to humans^5^. Mastitis is estimated to cost the global dairy industry up to 35 billion USD per year^6^.

Upon infection, *S. aureus* responds to environmental cues by adjusting expression of virulence determinants including secreted and envelope-associated molecules^7^, major metabolic reprogramming, and cell envelope remodeling^8,9^. Numerous systematic studies generated inventories of factors whose expression is altered in milk and serum^10,11^. However, the bacterial state, and the specific fitness factors associated with growth, survival and antibiotic adaptation in these infection-relevant environments have received little attention.

The Gram-positive bacterial envelope comprises a complex semipermeable macromolecular layer that optimizes bacterial fitness by sensing and responding to environmental signals. Fatty acids (FAs), the building blocks of membrane phospholipids, play a significant role in staphylococcal survival and adaptation to variable food and animal environments. Membrane FA composition impacts membrane fluidity, energy production, and virulence regulation, which determine how well bacteria withstand environmental stresses^12-14^. The *S. aureus* FA synthesis pathway (FASII) produces saturated branched-chain and straight-chain FAs^15^. In addition, *S. aureus,* like most known bacteria, scavenges environmental/host FAs, and incorporates them into its membrane phospholipids^16-20^. It is notable that these dynamic adjustments in response to exogenous FAs impact membrane properties and modulate bacterial function and survival to external stresses^21,22^. *S. aureus* also synthesizes the carotenoid pigment staphyloxanthin, which is embedded in the membrane *via* its rigid, planar structure and FA and isoprenyl lipid tails. Staphyloxanthin reinforces membrane structural integrity by reducing fluidity and favoring lipid packing. It is also implicated in membrane protein organization. Increased membrane rigidity may enhance its mechanical strength, and explain greater tolerance to environmental and host immune-mediated stress^20,23-25^.

*S. aureus* is notorious for its adaptation and virulence capacities in numerous biotopes. Here, we investigated *S. aureus* phenotypic properties in two environments relevant to infection, milk and serum. We report that both these environments modify *S. aureus* envelope and membrane properties, and protein production, but not in the same way. Unexpectedly, compared to serum, milk adaptation confers a marked fitness advantage, with a higher infectivity *in vivo*. These results have direct implications for *S. aureus* human and livestock mastitis, and for septic infection.

## Results

### *S. aureus* grown in milk or serum preferentially incorporate exogenous FAs in membrane phospholipids

Growth kinetics of MRSA derivative JE2 was first compared in laboratory (BHI) medium, milk, and serum over a 24h period. *S. aureus* grew robustly in 100 % raw cow milk or in BHI, with no visible latency phase (Fig. 1a). In contrast, MRSA growth in 100% adult bovine serum was preceded by a pronounced latency phase. Delayed growth might be due to inhibitors and/or stress conditions in bovine serum, e.g., linked to immunoglobulins, poor iron availability, and the presence of polyunsaturated FAs that exert antimicrobial activity against staphylococci^26^. In contrast to the observed multiplication of *S. aureus* in bovine serum, *S. aureus* reportedly survived but did not replicate in human serum^10^. In our conditions, growth initiated after six hours, i.e., beyond the usual test period; this, and/or species-specific serum variations may underly these differences. *S. aureus* cells grown and adapted to the two media will be referred to as serum-adapted and milk-adapted cells.

**Figure 1.**
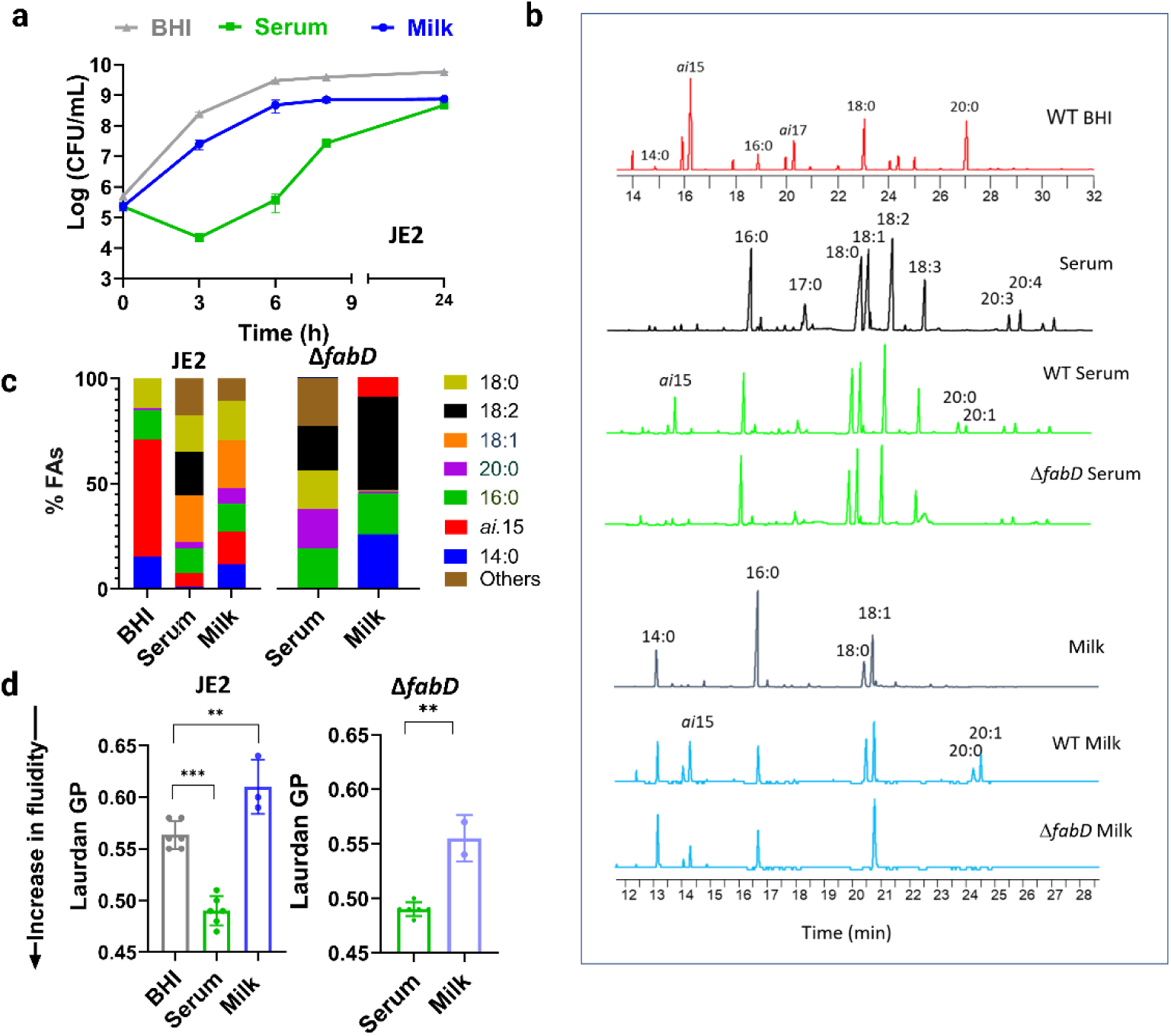
*S. aureus* growth and exogenous FA incorporation in serum or milk. (**a**) *S. aureus* JE2 growth was assessed by determining CFUs in serum, milk, and BHI liquid medium by plating dilutions on BHI agar at given time points. Data represent mean ± SD of three independent experiments. (**b**) Membrane FA profiles were compared in JE2 (WT) and the Δ*fabD* FA auxotrophic derivative grown in serum, milk, or BHI. The JE2 strain cultured in BHI (red profile) synthesizes FAs via FASII, and its FA profile is dominated by *ai*15, its elongation products (*ai*17, *ai*19) and saturated straight chain FAs (C16:0, C18:0, C20:0). In contrast, JE2 FA profiles in serum (green) or milk (blue), are both enriched with exogenous FAs supplied by the medium. The FA auxotroph Δ*fabD* displays a similar FA profile as that of its respective medium. Peak heights correspond to relative responses (mV) of each FA in a sample. Major FAs are indicated; N=3. (**c**) FA species are presented as the proportions of the total adjusted to 100%. Shown is the average of three independent experiments. See Table 1 in Supplementary Information for proportions of each FA per sample. (**d**) Membrane fluidity of WT JE2 and its Δ*fabD* derivative adapted to various media was evaluated using the Laurdan generalized polarization method (Laurdan GP). Laurdan GP = (I_440_ − I_490_)/(I_440_ + I_490_), where I_440_ and I_490_ are the emission intensities at 440 and 490 nm, respectively, when excited at 350 nm. Individual data points (n = 5 biologically independent samples) are shown together with mean ± SD. P values were determined by one-way ANOVA (*** p<0.0001; ** p< 0.0008).

Numerous bacteria including *S. aureus* incorporate environmental FAs directly in membrane phospholipids ^16,20,27^. We analyzed exogenous FA incorporation upon *S. aureus* growth in serum and milk by comparing their FA profiles with that of bacteria grown in BHI, which contains only trace amounts of FA. *S. aureus* FA profiles displayed differential incorporation of medium-derived FAs (Fig. 2b). BHI-grown *S. aureus* produced FAs corresponding to those synthesized by FASII, i.e., anteiso branched-chain FA *ai*15 and its elongation products *ai*17, and straight-chain FAs C18:0 and C20:0 (Fig. 1b), as reported^15,20,28^. By contrast, *ai*15 was a minor species in *S. aureus* grown in either bovine serum or milk (Figs. 1b and 1c). We also compared FA profiles of the JE2 strain with those of an isogenic Δ*fabD* mutant grown in serum and milk under similar conditions. FASII activity is turned off in strains lacking a functional *fabD* gene, which encodes the FASII initiation FabD protein (malonyl CoA-acyl carrier protein transacylase)^28^ (Supplementary Fig. 1). The Δ*fabD* strain, which relies on exogenous FAs, was able to replicate in serum and milk confirming that *S. aureus* can efficiently incorporate FA from these two media (Supplementary Fig. 2). Moreover, the FA profiles of both wild type (WT) and Δ*fabD* strains reflected the FA composition of the medium in which they were adapted (Fig. 1b).

**Figure 2.**
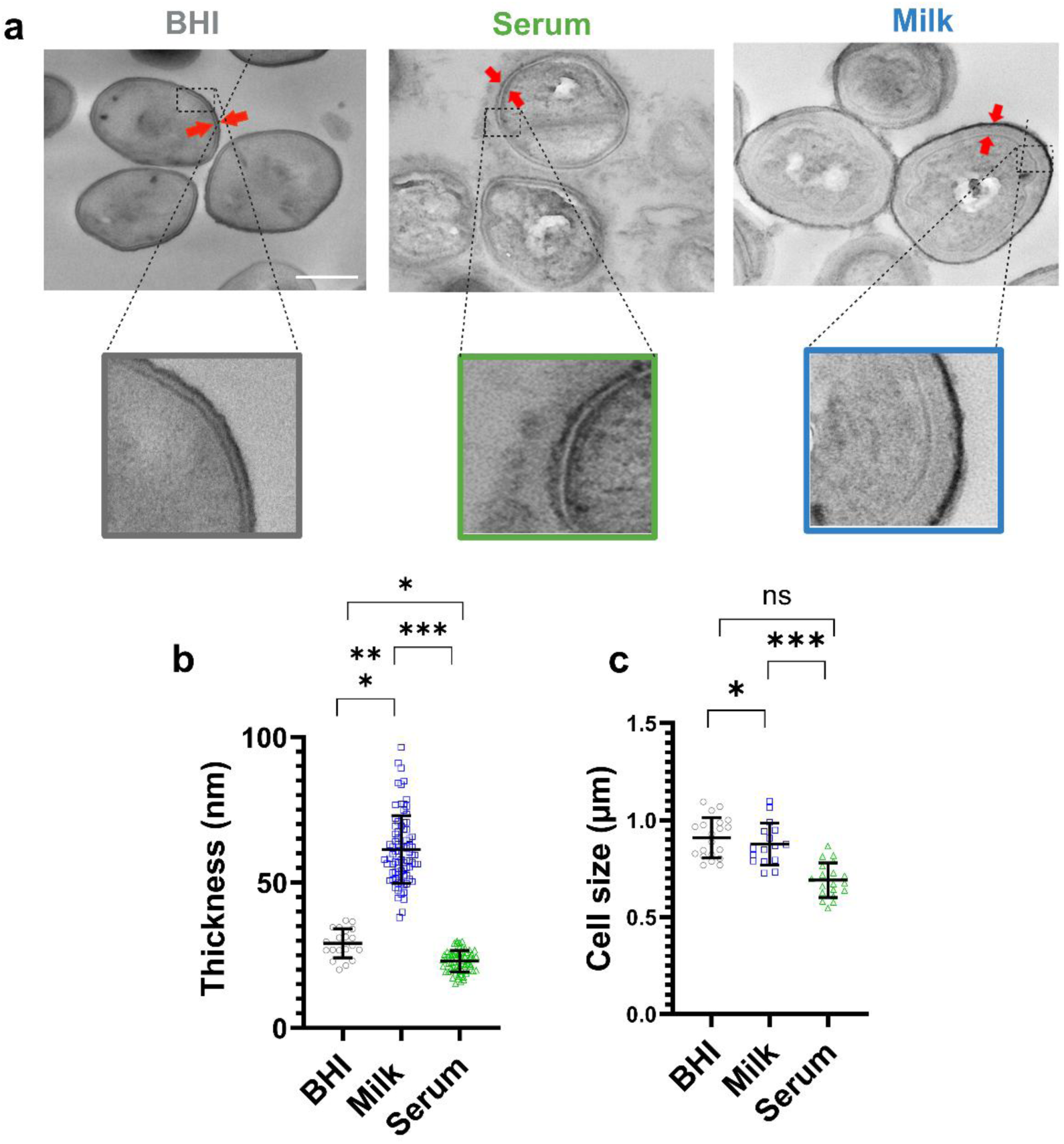
Serum and milk modulate *S. aureus* envelope. (**a**) Representative transmission electron microscopy images of milk-adapted and serum-adapted saturated JE2 bacteria. BHI-grown cells are presented for comparison. Arrows indicate envelope thickness. Scale bar for the upper panels, 500 nm. The zoom illustrates the highly different *S. aureus* envelope morphology between the 3 conditions. (**b**) Range of cell envelope thickness, and (**c**) cell sizes of JE2 adapted to BHI, serum, or milk, were determined on micrograph cross-sections using ImageJ. Each circle represents a measurement for a single cell. Boxplots of panels show the median diameter distribution (thick line within boxes) and the degree of variability (amplitude of the box). Error bars indicate SD. Statistical significance was determined by one-way ANOVA (*** p<0.0001; *, p<0.0160).

The WT *S. aureus* FA profile comprised both self-synthesized FA species and C18:1, which is necessarily incorporated from the medium as *S. aureus* does not synthesize unsaturated FAs (Supplementary Table 1 and Fig. 1c). Additional FA elongation products appeared in the WT strain, but not in Δ*fabD* whose FASII activity is disabled (Fig. 1b). Although *ai*15 comprises < 3% of total FAs in cow milk^29^ and was not detected in the milk extract, it represented 10% of FA present in the Δ*fabD* mutant. Surprisingly, no C18:0 was detected in the *ΔfabD* mutant adapted to milk, suggesting a bias towards incorporation of short chain saturated FAs, possibly including trace amounts of *ai*15 (Fig. 1c). The serum FA profile is more complex than that of milk, and in particular comprised long-chain unsaturated and polyunsaturated FAs (C18:1, C18:2, C18:3, C20:3 and C20:4) (Fig. 1b, middle profile). These FAs were incorporated in both Δ*fabD* and JE2 strains (Fig. 1b and 1c, and Supplementary Table 1). Elongated FA products were proportionately greater in milk-than in serum-adapted bacteria (Fig. 1b and 1c). Serum is rich in polyunsaturated FAs, which inhibit growth and the FASII pathway^30^.

While serum and milk are both lipid-rich, their FA compositions differ, and *S. aureus* displays a different FA profile reflecting the lipid resources of each biotope. Consequently, FA profiles of WT and Δ*fabD* (FA auxotroph) strains were similar for a given medium, confirming that in these environments, *S. aureus* preferentially incorporates exogenous FAs rather than synthesizing them *de novo*. The markedly different FA profiles of the same strain cultivated in different environments (JE2 profiles in Fig. 1b and 1c) led us to speculate that bacterial phenotypes would be impacted.

### Serum and milk have inverse effects on membrane fluidity

We considered that greater proportions of polyunsaturated FAs and overall lower FASII-mediated elongation in serum may impact membrane fluidity. We assessed membrane fluidity of milk-, serum-, and BHI-adapted *S. aureus* JE2 using the membrane fluidity-sensitive dye Laurdan^31^. Laurdan intercalates into the membrane bilayer and exhibits a fluorescence emission wavelength shift dependent on the amount of adjacent water molecules^31^, which depends on the packing density and fluidity of the lipid bilayer. Serum-adapted bacteria showed the greatest membrane fluidity, followed by BHI, while milk-adapted JE2 showed significantly lower membrane fluidity (Fig. 1d). The difference associated with serum- and milk-adapted conditions was also seen in the Δ*fabD* mutant (Fig. 1d). Increased membrane fluidity of serum-adapted *S. aureus* is consistent with the greater proportion of polyunsaturated FAs C18:2 and C18:3 (Fig. 1b and c), and is in line with looser lipid packing. In contrast, proportionately greater long-chain FAs in milk-adapted *S. aureus* cultures would favor greater membrane rigidity. Other membrane structures may also be contributing factors (see below). Collectively, these findings indicate that *S. aureus* membrane composition and fluidity are dynamically and differentially affected by FA-rich environments. In both media, *S. aureus* membrane FAs are dominated by exogenous FAs, albeit under very different physiological conditions. Milk appears to provide a more favorable environment for *S. aureus* growth than serum; the absence of potentially toxic polyunsaturated FAs in milk may be one reason for better growth.

### *S. aureus* envelope thickness, permeability, surface charge, and hydrophobicity vary between serum, milk, and BHI environments

The extreme differences in membrane FA composition, lipid packing density and membrane fluidity, suggested that other physical and functional *S. aureus* properties might vary between serum, milk, and BHI cultures. Transmission electron microscopy performed on ultrathin cross-sections of *S. aureus* stationary phase cells revealed well-separated spherical cocci (Fig. 2a). Envelopes of the milk-adapted *S. aureus* JE2 were markedly thicker than those of serum- or BHI-grown cells (Fig. 2a and Supplementary Fig. 3). Quantitative analysis of envelope thickness, excluding the thread-like structures around serum-adapted cells, confirmed that envelopes of milk-adapted JE2 were about 2-fold thicker than those of serum- or BHI-grown bacteria (39.4 nm compared to 22-29 nm respectively, Fig. 2b). Moreover, cross-sectional quantification of cell outer diameters showed that serum-adapted *S. aureus* were smaller compared to the two other groups (0.69 µm compared to 0.87-0.89 µm respectively, Fig. 2c). Despite the significantly thicker envelopes of milk-adapted cells compared to BHI-grown cells, their outer cell diameters were comparable.

Serum-adapted JE2 displayed irregularly textured surfaces compared to the smooth surfaces of milk- and BHI-grown bacteria (Fig. 2a and Supplementary Fig. 3). We asked whether the textured surfaces corresponded to peptidoglycan fragments on the cell surface, as reported in human serum^10,32^. Bodipy-Vancomycin, a cell wall-specific probe that binds to the D-Ala-D-Ala moiety, was used to evaluate differences in peptidoglycan staining in serum-, milk-, or BHI-grown *S. aureus* (Supplementary Fig. 4). Vancomycin binding was significantly greater in bacteria grown in serum compared to those grown in BHI (as shown previously^32,33^) or in milk (Fig. 3a). Markedly lower vancomycin binding in milk-adapted bacteria may be due, at least in part, to impeded antibiotic penetration through the thickened cell envelope, a phenomenon that can contribute to resistance in *S. aureus*^34^.

**Figure 3.**
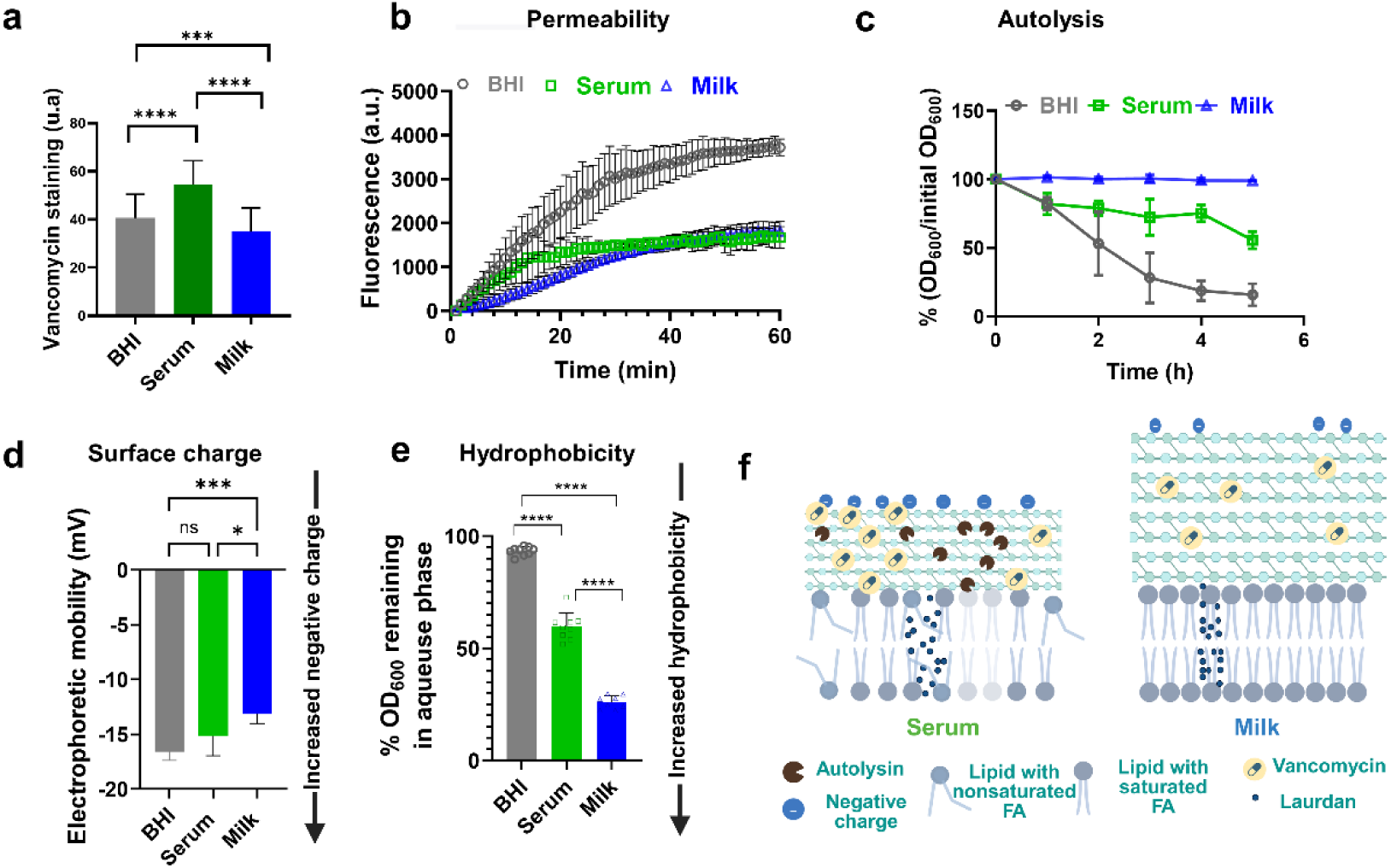
*S. aureus* physical features are differentially modulated by serum and milk. JE2 cells prepared in BHI, serum, or milk conditions washed and resuspended in PBS, were assessed for envelope properties. (**a**) Peptidoglycan accessibility was determined using an exogenous fluorescent probe, BODIPY^™^ FL-vancomycin (3 µg/mL). Fluorescence at 488 nm was measured and quantified using ImageJ. Graphs represent the mean ± SD of 3 independent experiments per condition. (**b**) Bacterial permeability was assessed by the EtBr diffusion assay^35^. Treatment with EtBr, which emits fluorescence upon internalization, was followed in bacteria incubated at 37°C. Fluorescence readings (excitation and emission wavelengths were 530 and 600 nm, respectively) were taken at 1-min intervals over a 60 min period. Values represent means of 3 biologically independent samples. (**c**) Sensitivity to autolysis was compared in cells resuspended in PBS with 0.5% Triton X-100 at time 0. Data presents mean ± SD of *n* = 3–4 biologically independent samples. (**d**) Cell surface charge was measured by electrophoretic mobilities, determined by zeta potential in bacterial cultures suspended in PBS. Data represent 5 independent runs with 3 replicates per run and are presented as mean ± SD. (**e**) Hydrophobicity was determined based on the bacterial affinity between PBS and xylene solvent. The affinity to adhere to liquid hydrocarbons was estimated by OD_600_ measurements in the aqueous phase before and after mixing. Results show the average and range of biologically independent triplicates. P values were determined by one-way ANOVA (**** p<0.00001; *** p<0.0001; ** p< 0.0008; * p=0.0160). (**f**) Schematic summary of *S. aureus* envelope specificities comparing milk and serum biotopes based on the physical features demonstrated above. Source data are provided as a Source Data file.

*S. aureus* permeability, autolysis, surface charge, and hydrophobicity were assessed and compared as a function of the respective growth environments. Permeability was analyzed using the ethidium bromide (EtBr) diffusion assay which also indirectly measures efflux^35^. The thick peptidoglycan layer of Gram-positive bacteria permits small molecules like EtBr to penetrate via passive diffusion, where it intercalates with DNA, enhancing fluorescence emission. Compared to bacteria grown in BHI, both serum- and milk-adapted cells showed lower permeability values (Fig. 3b). Low permeability of EtBr suggests greater resistance of bacteria grown in natural media to the diffusion of small molecules in and out of bacteria, which could in turn modify bacterial susceptibility to exogenous drugs. However, since permeability was similar in serum- and milk-adapted bacteria, this characteristic would not account for differences between the two conditions.

JE2 sensitivity to autolysis in the serum-, milk-, and BHI-adapted cultures was assessed using a Triton X-100 based assay^35^. Autolysis of BHI-grown bacteria was nearly complete at 5 h post-treatment, while that of serum-adapted bacteria was less pronounced (15 % and 55.5 % of the initial optical density, respectively). Strikingly, no autolysis was detected in milk-adapted bacteria (Fig. 3c). This correlates with the significantly thicker cell envelopes observed by electron microcopy (Fig. 2a) and might suggest that *S. aureus* autolysin activity is lower in milk. Weakly lytic bacteria are associated with increased virulence and resistance to antibiotics^36,37^.

Surface charge and hydrophobicity of JE2 were assessed according to growth medium by zeta potential measurements and affinity through a biphasic partitioning assay, respectively. The *S. aureus* electrostatic charge is conferred by numerous surface factors including membrane and cell wall proteins, ionized phosphoryl groups of the wall teichoic acid (WTA) and lipoteichoic acid (LTA), and unsubstituted carboxylate moieties of peptidoglycan that are exposed to the external environment^38^. Milk-grown bacteria displayed a less negatively charged surface than serum- or BHI-grown bacteria (Fig. 3d). Biphasic partitioning assay showed that milk-adapted JE2 cells were also significantly more hydrophobic than either serum- or BHI-adapted bacteria, such that milk>>serum>>BHI (Fig. 3e).

Taken together, milk and serum environments provoke discrete changes in the *S. aureus* envelope, likely through alterations in cell wall architecture and surface-associated molecules (Fig. 3f). Permeability was comparable in serum- and milk-adapted bacteria, suggesting that this property was not intrinsic to medium-specific adaptations. In contrast, milk-adapted *S. aureus* is distinguished by reductions in membrane fluidity, negative surface charge, autolysis, and vancomycin binding, but greater envelope thickness and hydrophobicity. These features may affect *S. aureus* fitness when adapted to a milk medium.

### Proteomic analysis in milk-adapted, serum-adapted, and BHI-grown *S. aureus*

We expected that the distinct morphological and physicochemical properties of *S. aureus* in serum and milk environments would correlate with altered bacterial protein expression. A label-free quantitative proteomics approach was used to identify *S. aureus* factors specific to the serum and milk biotopes, with BHI-grown bacteria as reference (Fig. 4a). A principal component analysis (PCA) based on imputed LFQ (label-free quantification) intensities for which proteins needed to be present in at least 50% of the samples in one group showed that the biological replicates clustered together, while hierarchical clustering analysis revealed significant variations in protein profiling between conditions (Supplementary Figs. 5 and 6). The proteome contained a total of 1355 proteins among the three groups, of which 44 and 41 were unique for serum- and milk-adapted cells, respectively; 57 proteins were detected in serum and milk, but not in BHI (Fig. 4b, Supplementary Data 1). In addition to the differentially detected proteins, some 600 proteins were enriched in a given category compared to the BHI-grown controls (using a p-value threshold of <0.05 and a fold change [FC] threshold of >1.5 or < 0.67) (Supplementary Fig. 6). A volcano plot revealed that in total 922 proteins were variable in serum- and milk-adapted conditions, of which 467 were significantly upregulated and 455 were significantly downregulated in serum-adapted vs. milk-adapted bacteria (P < 0.05) (Fig. 4c and Supplementary Data 1). Compared to the BHI-grown controls, 672 proteins were differentially expressed in milk (332 up-regulated and 340 down-regulated) and 642 in serum (443 upregulated and 209 downregulated proteins) (Supplementary Fig. 7).

**Figure 4.**
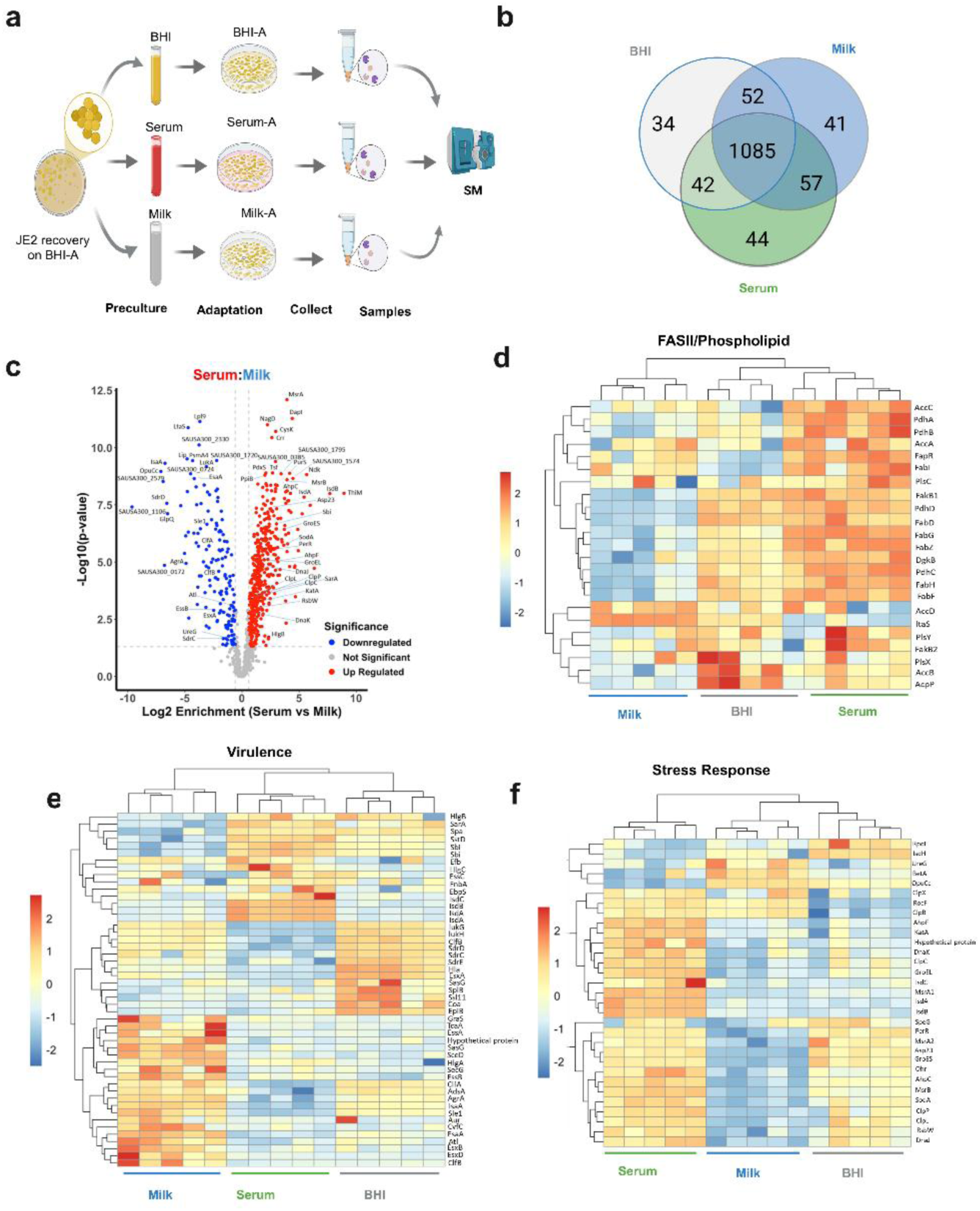
Comparative proteomics of BHI-, serum-, and milk-adapted *S. aureus*. (**a**) Flowchart for the study. Bacterial grown in the three conditions and in five independent replicates were harvested from agar plates after overnight cultures and processed for proteomics analysis. (**b**) Venn diagram showing the number of identified proteins detected in each condition, based on the five biological replicates. Full data is presented in Supplementary Data 1. (**c**) Volcano plot depicts the magnitude (log_2_ fold change) and statistical significance (q-value) comparing levels of the expressed proteins in JE2 between serum- and milk-adapted cultures. Horizontal dashed lines indicate fold-change threshold, vertical dashed lines indicate statistical significance threshold (q-value = 0.05). Red spots indicate significantly upregulated proteins, and blue spots indicate significantly downregulated proteins, comparing serum to milk cultures. See Supplementary Fig. 6 for volcano plot comparisons of serum *vs* BHI and of milk vs BHI. Heat maps of significantly different proteins (Z-scored) related to (**d**) the FASII pathway, (**e**) stress-response and (**f**) virulence, compare BHI-, serum-, and milk-adapted JE2. The expression level scale is shown at left. All five samples are shown for each condition.

We screened for membrane-related functions that might explain the observed morphological and physical differences of *S. aureus* in the 3 conditions. Among them, proteins related to lipid synthesis were most abundant in serum -adapted samples (Fig. 4d). This was initially unexpected as no FASII-mediated elongation of incorporated exogenous FA was observed in serum (Fig. 1b). We may speculate that poor growth in serum could stimulate FASII, while the overwhelming abundance of lipids present in milk (∼4% lipids) might suppress production of lipid synthesis-related proteins ^30^.

Lipidated molecules are common membrane components that contribute to bacterial interactions with the environment and membrane integrity. Lipoproteins are frequently receptors that sense the environment and are involved in metabolite and nutrient homeostasis^39^. Approximately half of the 36 detected putative lipoproteins were enriched in milk-adapted, compared to serum-adapted bacteria (Fig. 4d). Increased lipoprotein levels may reflect the active metabolism of milk-growing bacteria. Additionally, LTA synthase (LtaS) is required for synthesis of LTA, which contributes to bacterial envelope integrity and autolysin control, and is implicated in virulence^40,41^. LtaS levels were higher in milk-than in serum-adapted bacteria (Fig. 4d), which correlates with reduced autolysis in milk-adapted bacteria^42^ (Fig. 3c).

The physical features of milk-adapted *S. aureus* may be linked to other proteomic differences. The GraRS two component regulator is implicated in envelope thickness/autolysin resistance, and vancomycin tolerance^10^. Here, GraS (SAUSA300_0646) was only detected in milk-adapted bacteria and thus correlates with thickened cell wall and autolysis resistance phenotypes (Figs. 2, 3c, and 3d; Supplementary Table 2). Cell surface charge reportedly relates to two main structures: the negatively charged LTA and the positively charged lysyl phosphatidylglycerol^43^. LTA represents ∼12 Mole % of surface lipids^44^, but its charge is diminished by alanylation involving DltD poly-D-alanine transfer protein (SAUSA300_0838). DltD levels are highest in milk-adapted bacteria (Supplementary Data 2), which might lower global surface charge. In contrast, phosphatidylglycerol lysyltransferase MprF (SAUSA300_1255), which adds a positive charge to phospholipids, is less expressed in milk. Further studies will be needed to confirm the functions responsible for surface charge decreases in milk. Higher levels of GraS and DltD may contribute to these changes.

Unexpectedly, milk-but not serum-adaptation led to differential increases in virulence factors (Fig. 4e, Supplementary Fig. 8). They include six components of the Type VII secretion system (Supplementary Data 2); of note, the Type VII components are induced by FAs^45^, which are highly abundant in milk. The Cys-tRNA(Pro)/Cys-tRNA(Cys) deacylase efflux system and the multidrug efflux pump quinolone resistance protein NorB are uniquely expressed in milk- and serum-adapted cells, respectively. The efflux activity of these systems might contribute to the low diffusion of EtBr in the permeability test in both conditions (Fig. 3b). The iron-regulated surface determinants (Isd system), which acquire heme under nutrient limitation^46^, were upregulated in serum-adapted cells. This is consistent with known iron-deficiency of serum^26^ which could also lead to slow growth (Fig. 1a). Iron starvation may disrupt the electron transport chain leading to accumulation of reactive oxygen species (ROS)^47^. Indeed, damage repair and ROS proteins, such as AhpF, KatA, MsrA1, ClpC, and GroEL are up-regulated in serum-adapted cells (Fig. 4f), supporting this hypothesis. Furthermore, polyunsaturated FAs incorporated from serum are subject to oxidation and may stimulate production of oxidative repair proteins^48^. We can suppose that bacteria adapted in the hostile environment imposed by serum may be more vulnerable to further oxidative stress or antimicrobials. The significance of the distinguishing properties of milk- and serum-adapted *S. aureus* led us to examine bacterial tolerance to stress according to their environmental history.

### Intense pigment production by milk-adapted *S. aureus* contributes to stress resistance

The *S. aureus* pigment staphyloxanthin constitutes a first line of defense against oxidative stress^49,50^. Milk-grown *S. aureus* showed pronounced pigmentation (Fig. 5a). Spectral profiles confirmed the presence of a triple-peak characteristic of staphyloxanthin^50^ (Fig. 5b). Pigment was also detected in extracts of BHI-grown *S. aureus*. In contrast, pigment in serum-adapted cell extracts was below detection levels. Staphyloxanthin contributes to rigidifying the bacterial membrane^24,50,51^. The high degree of pigmentation in milk-grown *S. aureus* compared to serum also correlates with the marked differences in membrane fluidity in these conditions (Fig. 1d). Interestingly, greater pigmentation in milk-adapted *S. aureus* did not correlate with the proportion of membrane-derived *ai*15, the FA moiety of staphyloxanthin^52^. Possibly, low amounts of *ai*15 present in membrane phospholipids are sufficient for production of staphyloxanthin; alternatively, the FA moiety on staphyloxanthin might be variable when *ai*15 is unavailable.

**Fig. 5.**
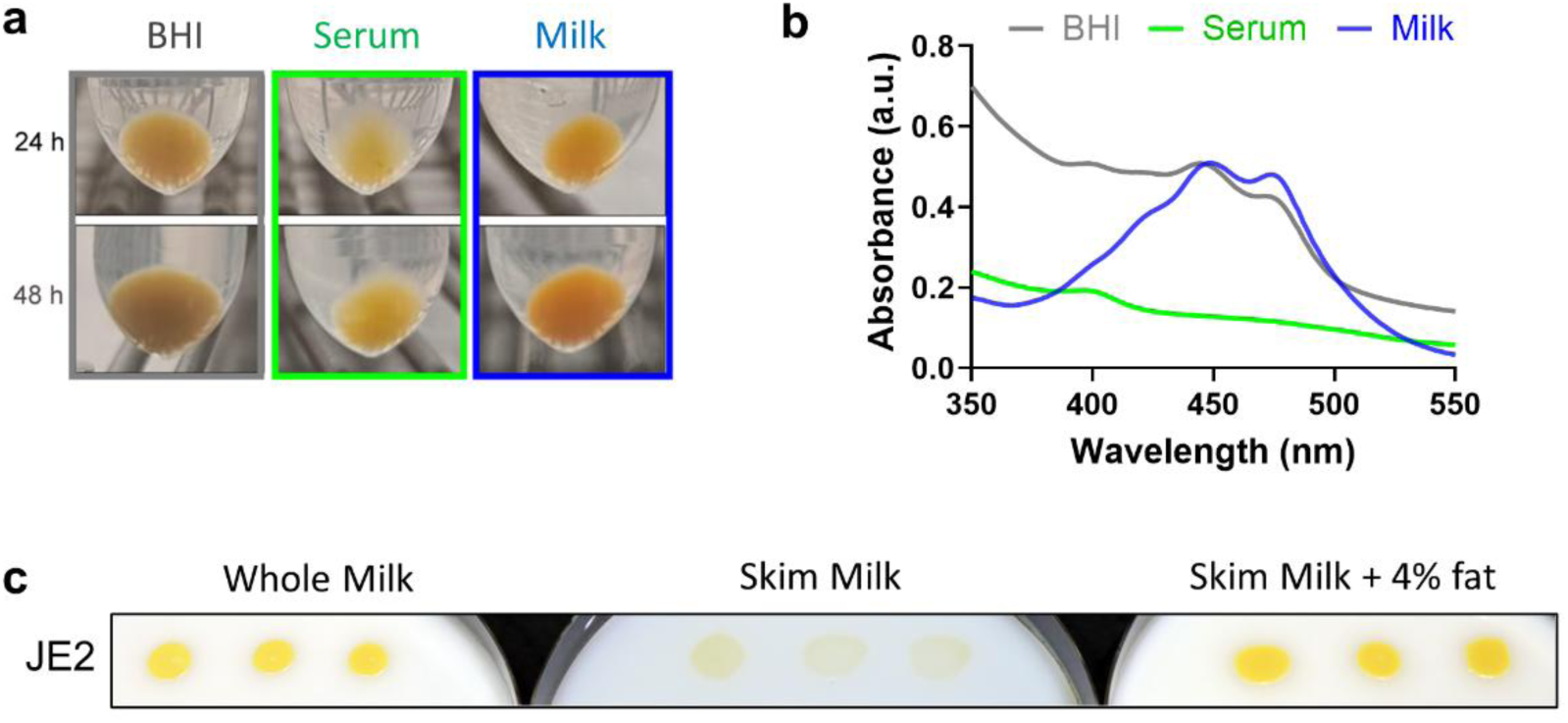
Milk-adapted *S. aureus* displays pronounced pigmentation that is stimulated by milk lipids. (**a**) Pigmentation was assessed in serum-, milk-, and BHI-adapted JE2 in cell pellets harvested after 24h and 48h growth. Cell pellets from 2 mL liquid cultures were collected, washed and photographed. (**b**) Pigment was extracted from cells and assessed by UV-Vis spectroscopy. A triple peak is due to the presence of the carotenoids staphyloxanthin, 4,4’-diaponeurosporene and 4,4’-diapolycopene, at ∼ 412 nm, 435 nm, and 465 nm respectively. Determination is representative of 3 experiments. (**c**) Independent JE2 cultures prepared in BHI were washed and adjusted to OD_600_ of 0.025 in 0.9% NaCl. For each culture, 5 µL were spotted on 1.5 % non-nutrient agar supplemented with 40% whole milk, skim milk, or skim milk supplemented with 4% milk cream. Plates were photographed after 48h.

As staphyloxanthin pigment comprises a lipid moiety^49^, we asked whether the milk lipids affect staphyloxanthin production. Plates containing whole milk, skim milk, or skim milk supplemented with milk cream (adjusted to 4% as present in whole milk) were spotted with the same JE2 cultures (Fig. 5c). Pigment was visibly lower at 48h on the skim milk plates. This result shows that lipids in the milk environment are required for high pigment expression.

The observed differences in pigmentation (Fig. 5a), membrane fluidity (Fig. 1d), and levels of stress response proteins (Fig. 4f) between milk- and serum-adapted *S. aureus* suggested that these growth conditions differentially impact bacterial susceptibility to oxidative and antimicrobial stress. Strikingly, treatment by singlet oxygen species (generated by 6 μg/ml methylene blue under visible light illumination) resulted in a 4-log greater resistance to killing in milk-adapted bacteria compared to serum-adapted cells (Fig. 6a). It was possible that milk components stayed bound to bacteria despite extensive washing, and interfered with the effects of ROS. To address this, serum-grown JE2 was incubated in raw milk for 1h (a time insufficient for regrowth), to enable milk molecules to adsorb, and were then washed and exposed to singlet oxygen. Post-incubation (pi) in milk had no effect on susceptibility of serum-grown *S. aureus* to singlet oxygen (Fig. 6a), indicating that the resistance phenotype observed in milk was directly due to the *S. aureus* cell state.

**Fig 6.**
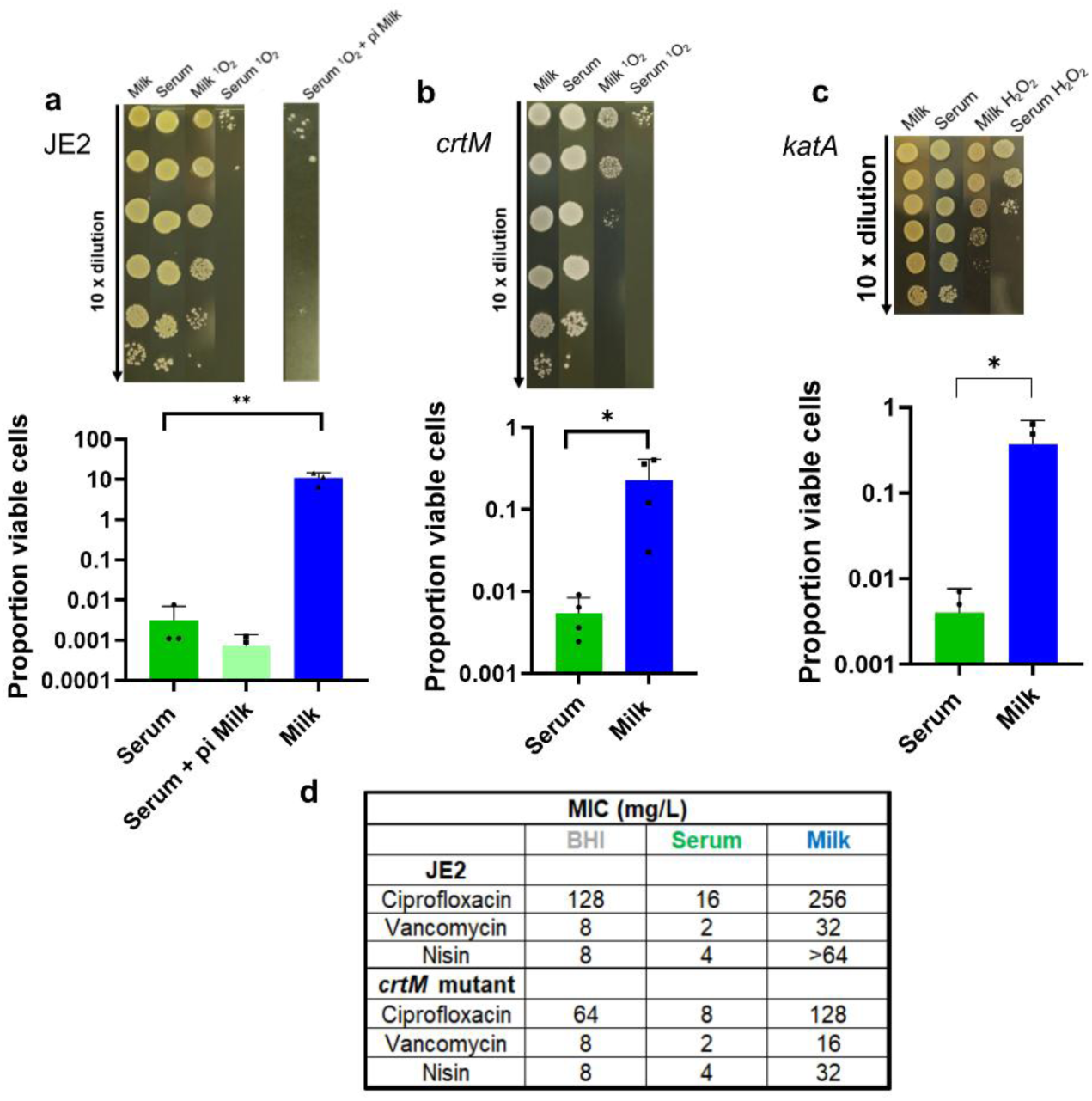
Tolerance to ROS and antibiotic is stimulated in milk-adapted *S. aureus*, and is diminished in *crtM* or *katA* mutants. JE2 WT and mutant cultures prepared in serum or milk conditions were washed and resuspended in physiological saline. (**a**) WT JE2 and (**b**) a *crtM* transposon mutant^75^ bacteria were untreated or exposed to singlet oxygen, and then serially diluted, spotted on BHI agar and grown overnight to determine survival. Experiments were repeated at least three times. For JE2 (**a**) an aliquot of serum-adapted bacteria was subject to 1 h exposure to milk and then washed prior to singlet oxygen treatment (at right). Continued high sensitivity allowed us to rule out non-specific effects of milk-bound bacteria. (**c**) H_2_O_2_ sensitivity was compared in milk-*versus* serum-adapted WT JE2 (Supplementary Fig. 10) and the *katA* transposon mutant^75^. For **a-c**, results are representative of at least 3 independent experiments. Upper panels show results of representative experiments. Note that all lanes within one image derive from simultaneously performed experiments, where different plates were used for determinations. Lower panels, error bars, SD. P values were determined by an ordinary one-way ANOVA (* p<0.016). (**d**) MIC, minimum inhibitory concentrations as determined by turbidity for different antibiotic concentrations after 24 h. Saturated bacterial cultures prepared in the indicated media were diluted in corresponding medium and dispatched in 96-wall plate. Bacteria were exposed to antibiotic concentrations in 2-fold dilutions ranging from 8 to 0.125 µg/mL (vancomycin), 0.5 to 0.0075 µg/mL (nisin) and (ciprofloxacin). Bacterial survival in milk was calculated after dilution spotting on BHI agar. Results are reported as mean of 3 independent experiments.

The role of staphyloxanthin in ROS tolerance was determined by testing a non-pigmented JE2 *crtM* transposon mutant (*crtM*::Tn) deficient in dehydrosqualene synthase (CrtM), which catalyzes the first step of staphyloxanthin biosynthesis^49^. The milk-adapted *crtM* mutant was more sensitive to singlet oxygen treatment compared to the parental strain (Fig. 6b). Nevertheless, compared to serum-adapted *crtM*, the milk-adapted *crtM* mutant still showed 1-2-log greater survival to singlet oxygen treatment, indicating that factors other than staphyloxanthin contribute to ROS tolerance. Similarly, a non-staphyloxanthin-producing strain, RN-R (a *S. aureus* RN4220 derivative repaired for the *fakB1* gene^53^), showed intermediate ROS survival in milk, while RN-R cells adapted to serum were eliminated (Supplementary Fig. 9).

The host produces hydrogen peroxide (H_2_O_2_) in response to infection^54^. We compared survival of milk- and serum-grown JE2 exposed to 1.5% H_2_O_2_ in PBS. However, no killing was observed in any bacteria in these conditions (Supplementary Fig.10). Given that *S. aureus* produces a catalase that metabolizes H_2_O_2_ into O_2_ and H_2_O^55^, we tested the JE2 *katA* transposon mutant (*katA*::Tn), a catalase-deficient strain, for H_2_O_2_ sensitivity. The milk-adapted *katA* mutant showed a 1-2-log greater tolerance to H_2_O_2_ than the serum-adapted strain (Fig. 6c).

Together, these results show that milk-adapted *S. aureus* have a marked survival advantage against oxidative stress over serum-adapted cells. Greater staphyloxanthin production in milk, which is stimulated by lipids, confers the major survival advantage. Overall, the comparatively increased sensitivity of serum-adapted *S. aureus* to oxidative stress compared to that in milk (Figs. 6a-c) indicates that increased levels of stress response proteins may not confer greater bacterial fitness.

### Milk growth increases *S. aureus* phenotypic resistance to front-line antibiotics and a bacteriocin

Numerous studies associate oxidative stress adaptation to antibiotic resistance^5,9,56^. High oxidative stress tolerance of milk-adapted *S. aureus* led us to ask whether sensitivity to ciprofloxacin, vancomycin, or nisin was also altered in the milk biotope. Ciprofloxacin inhibits DNA replication by blocking gyrase and topoisomerase, and induces oxidative stress^57,58^. ROS scavengers, such as ascorbic acid, exacerbate ciprofloxacin antibacterial activity^59^. Vancomycin, a last-resort antibiotic to treat MRSA infections, targets the D-Ala-D-Ala moiety of peptidoglycan precursors to inhibit peptidoglycan crosslinking; it also binds to lipid II^58^. Nisin, a bacteriocin primarily used as a food preservative, docks to peptidoglycan precursor lipid II to permeabilize the membrane and inhibit cell wall synthesis^58^.

Serum- or milk-adapted *S. aureus* were washed and tested for antimicrobial sensitivity by a dilution assay. Serum-adapted cultures were the most susceptible to all three antimicrobial treatments, while milk adaptation conferred the highest resistance (Fig. 6d). Antibiotic sensitivity was augmented in *crtM* mutants, particularly in milk-adapted conditions (Fig. 6d).

Greater antimicrobial resistance in milk may be multifactorial: (i) ROS exacerbates ciprofloxacin, but not vancomycin or nisin sensitivity. This was confirmed by measuring antibiotic inhibition zones without or with ascorbic acid addition (Supplementary Fig. 11). Ciprofloxacin-generated ROS causes more damage in the *crtM* mutant, which shows accrued ROS sensitivity compared to the WT strain (Fig. 6a). (ii) Greater membrane rigidity and a thicker envelope of milk-adapted bacteria (Figs. 2 and 3a) might reduce nisin and vancomycin accessibility to D-Ala-D-Ala moieties. In addition, lower levels of the regulator SarA (SAUSA300_0605) in milk-adapted bacteria (Fig. 4c, Supplementary Data 1) correlates with vancomycin resistance^60^. (iii) Interactions of positively charged antimicrobials like nisin with bacteria may be affected by surface charge; reduced negative charge in milk-adapted *S. aureus* (Fig. 3d) might lower nisin binding. These considerations suggest that various properties of milk-adapted *S. aureus* led to phenotypic antimicrobial resistance. The specific contributions of these properties needs to be examined in future work. The above data illustrate the importance of the environment for determining stress resilience and antimicrobial tolerance of bacteria. Greater fitness of *S. aureus* adapted to milk compared to serum indicates that a single bacterium may have widely varied fitness states and that milk-adapted bacteria are well equipped to survive hostile conditions during infection. They also suggest that mastitis infections in mammals carrying milk might require specific approaches for treatment.

### Milk-adapted *S. aureus* has increased virulence in an insect larvae model

We used a *Galleria mellonella* infection model to assess how bacterial history, i.e., *S. aureus* issued from BHI, serum, or milk adaptation, affects insect mortality (Fig. 7). Upon infection, *G. mellonella* produces antimicrobial peptides and ROS that accumulate in the larval haemocoel, the fluid analogous to blood^61^. These properties and the use of large cohorts make *G. mellonella* a suitable model to obtain a statistically significant assessment of *S. aureus* virulence in a simplified system. Larva (60 per cohort) were injected with 10^6^ CFU BHI-, serum-, or milk-adapted *S. aureus* JE2 directly into the hemocoel, or with physiological saline, and mortality was recorded daily for 3 days (Fig. 7a). Virulence was defined as increased larva mortality due to infection. Insects infected with milk-adapted JE2 showed the greatest mortality at 24 h compared to serum- or BHI-grown bacteria (Fig. 7b). These differences diminished after 48 h, suggesting that the bacterial advantage by its previous environment is transient. We conclude that adaptation to natural environments can shape *S. aureus* virulence properties *in vivo*.

**Figure 7.**
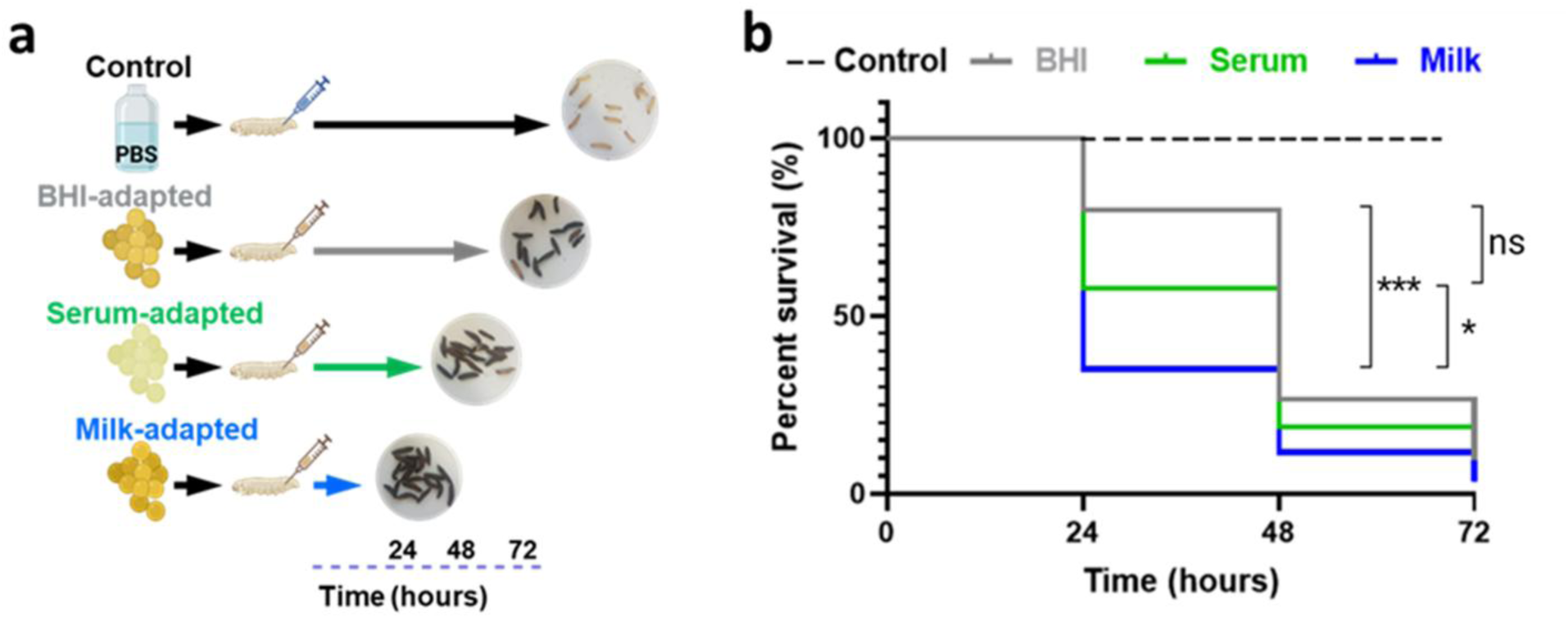
The *S. aureus* environmental origin affects infection efficacy. The infection capacity of serum-, milk-, or BHI-adapted *S. aureus* was compared using the *G. mellonella* infection model. (**a**) Schematics of the *In vivo* infection assay. Larvae were infected with 10^6^ CFUs of *S. aureus* JE2 issued from the 3 conditions. Experimental control groups received PBS. Insect killing was assessed by melanization. (**b**) Surviving larvae were counted after 24h, 48h and 72h (n=20 larvae/group; 3 independent experiments); insect survival in PBS treated insects showed no mortality. Data were analyzed using Kaplan-Meier with pooled values of biologically independent triplicates (60 samples per condition). The log-rank (Mantel-Cox) test was used to compare survival curves. Significances between the 3 groups is indicated: *** p<0.001; * p>0.05; ns: not significant.

## Discussion

Bacterial adaptation is essential for survival in changing environments. The present work reports that *S. aureus* exposure to different environments causes specific structural and expression phenotypes that shape envelope physical properties, membrane composition and rigidity, and tune fitness functions. Milk-adapted *S. aureus* is characterized by extreme ROS resistance associated with pronounced pigmentation, and phenotypic antibiotic resistance. The shift towards greater virulence factor production in the milk environment supports the hypothesis that bacteria can be primed for infection, as confirmed in an insect infection model.

### Physiological rationale for *S. aureus* fitness in milk in the context of infection

The physical features and specificities in protein expression and pigment production of milk*-*adapted *S. aureus* explain their greater fitness in withstanding oxidative and antibiotic stress, and accelerated virulence in the insect infection model compared to serum-adapted bacteria. These observations are consistent with *in vivo* conditions, where *S. aureus* CFUs are overall low or undetectable in blood but high in organs.

It was initially perplexing that *S. aureus* virulence potential is activated in a ‘food source’ and is superior to that in serum. However, *S. aureus* is among the primary causes of mastitis in livestock, where udders contain infected milk^62^. *S. aureus* that contaminates milk is thus part of the infectious process, and increased bacterial fitness would exacerbate infection. While USA300 was not implicated in mastitis in cattle, this lineage was reported to cause breast infections during parturition in women^63^. Milk is thus not only a food biotope, but also a possible host reservoir and infection portal.

Since numerous stress response protein levels were significantly higher in serum, we also expected that adaptation to serum would confer an advantage for stress tolerance compared to milk (Figs. 6a-c). However, the opposite was observed, i.e., oxidative stress tolerance was markedly diminished compared to that in milk, suggesting that increased levels of stress response proteins may not confer greater bacterial fitness. Reduced fitness in serum despite the induction of numerous stress-defense factors reflects the *in vivo* situation, where *S. aureus* CFUs are generally low or undetectable in blood as compared to organs^64^. We note that serum composition varies between species and individuals, which may modulate the properties reported here in bovine serum, and in human serum studies elsewhere^10,33^. The use of pooled bovine serum and milk sources gives confidence to the reported properties in each biotope. As a first comprehensive comparative study, it demonstrates that these two biotopes cause a single bacterium to adapt via very different properties.

### A role for membrane lipids in dictating fitness and virulence

Intense *S. aureus* pigmentation in milk contributes to greater fitness, in particular to oxidative stress (Figs. 6a-c). We speculate that most fitness advantages of milk-compared to serum-adapted *S. aureus* relate to environmental lipids whose FA moieties are incorporated into bacterial membranes. Milk contains about 10-fold more lipids than serum, which are mainly triglycerides that supply incorporable FAs^65,66^. Bacteria grow similarly in whole and skim (fat-free) milk^67^, indicating that milk lipids are not toxic despite their high concentrations. Both milk and serum lipids markedly alter *S. aureus* membrane FA composition, but not in the same way. In both media, FASII-synthesized *ai*15 and its elongation products are decreased (Fig. 1c). Lipids from serum but not from milk comprise polyunsaturated FAs, whose incorporation in *S. aureus* membranes increases membrane fluidity (Fig. 1d), sensitizes bacteria to ROS damage^12,48^, and is associated with antibiotic sensitivity (Fig. 6 a-d).

*S. aureus* incorporation of milk lipids is required for increased staphyloxanthin pigment formation (Fig. 5c). Interestingly, a failure to incorporate environmental FAs in a *S. aureus fak* (encoding fatty acid kinase) mutant causes a defect in pigment production^68^, supporting the role of environmental lipid incorporation in stimulating pigment formation. Pigment contains lipid moieties, and its production in milk was the main factor involved in ROS resistance (Figs. 6a-c). This suggests that the gross advantage of growth in milk relies on changes in *S. aureus* membrane lipids (pigment is one of them). A study of milk-induced effects on *S. aureus* expression, published in the course of our work, highlighted metabolic factors whose requirements were alleviated in milk^11^. The 8325-4 lineage used in that report is defective for FA incorporation due to a *fak* defect, and fails to produce pigment^53,68^; therefore, the effects of milk lipid incorporation on *S. aureus* fitness as reported here would have gone undetected. Since 95% of *S. aureus* carry intact *fak* genes, we consider that a USA300 derivative is representative of livestock or clinical strains. Combining information from these studies discriminates between the effects of milk that rely on host lipids from those that do not. Based on these comparisons and our findings, we conclude that the major *S. aureus* fitness advantages upon milk adaptation are related to membrane lipid alterations, while the physical barrier related to envelope thickening in milk-adapted bacteria likely contributes to decreased antibiotic sensitivity to vancomycin and nisin.

## Methods

### Bacterial strains, media and growth conditions

*S. aureus* strains used in this study are listed in Supplementary Table 2. Brain heart infusion (BHI) is the base medium of *S. aureus* cultures unless specified. The *ΔfabD* mutant was grown in BHI containing the appropriate FA mixture (C14:0, myristic acid; C16:0, palmitic acid; and C18:1*cis*, oleic acid (Larodan Fine Chemicals, Sweden), each added to a final concentration of 170 µM from 100 mM stocks prepared in DMSO) and adult bovine serum (from cattle older than 12 months, Eurobio, France) at 10 % final concentration, and is referred to as SerFA medium. Bacteria were also cultured in commercial organic pasteurized whole milk, skim milk (Lactel, France), and skim milk containing 30% fat in cream (Franprix, France). Solid plates were prepared by adding 20% or 40% of milk products as specified, or 20% of the adult bovine serum to 1.5 % non-nutrient agar (Invitrogen, France) in H_2_O. *S. aureus* was cultured in liquid media in aerobic conditions at 37°C for 18 h to reach stationary phase. To monitor growth kinetics *S. aureus* overnight cultures were diluted to an OD_600_ of 0.025 and grown in different media at 37°C with shaking. Growth was monitored by measuring OD_600_ every 30 min or by plating dilutions on BHI agar. Growth curves were performed based on at least three biologically independent samples.

### Construction of a *S. aureus* USA300 *fabD* deletion mutant

plsX and fabG (primers for plsX fragment were designed based on the JE2 DNA sequence: pMad-plsX_F and plsX_fabG_R; primers for fabG fragment: plsX_fabG_F and fabG_pMad_R fwd; see Supplementary Table 3) were PCR-amplified from USA300-JE2 genomic DNA. The pMad thermosensitive vector ^69^ was digested with SmaI (NEB R0141). The three fragments were assembled according to the one-step isothermal DNA assembly by the Gibson protocol (NEB – Gibson Assembly Cloning Kit, France). The mixture was incubated at 50°C for 30 min in a thermocycler (Mastercycler, Eppendorf, France). The ligation mix was used to transform 100 µL chemo-competent *E. coli* DH5α cells and plated on LB agar with 100 µg/mL ampicillin. PCR on colonies was performed with pMad oligonucleotides to identify clones. Bacterial plasmid DNA extraction was performed using QIAprep Spin Miniprep Kit (Qiagen). The pMad-plsX-ΔfabD-fabG plasmid isolated from *E. coli* was introduced by electroporation into *S. aureus* RN4220-R (a RN4220 derivative in which the *fakB1* defect was corrected^53^) and transformants were selected at the permissive temperature (30°C) for resistance to erythromycin (5 µg/mL). Plasmid pMad-plsX-ΔfabD-fabG was extracted from RN4220-R with a preliminary lysis treatment of *S. aureus* cell suspensions with lysostaphin (100 µg/mL) for 1 h at 37°C, and was used to transform electrocompetent *S. aureus* JE2. Chromosomal *fabD* inactivation was achieved as described^70^, with the following modifications: an overnight preculture was grown at 30°C in BHI medium with 5 µg/mL ery and diluted 100-fold in the same medium with selection. After 2 h growth at 30°C, the culture was shifted to the nonpermissive temperature at 42°C for 6h to permit the plasmid integration into the *S. aureus* chromosome by homologous recombination. As *fabD* is essential for *S. aureus* FA synthesis, exogenous FAs were provided, and mutant candidates were grown at 42°C on SerFA plates with erythromycin. To select for the second recombination event, an overnight culture at 42°C of the single crossover mutant was grown with erythromycin (3 µg/mL) in SerFA liquid medium, diluted 1/1000-fold in SerFA medium with no selection, and then maintained at 30°C for 6 h to favor plasmid excision. Cultures were plated at 42°C on SerFA for 30 h-48 h. Clones were verified for *fabD* gene inactivation phenotypes by PCR on colony and sequencing.

### Determination of *S. aureus* FA profiles

FA profiles were determined as described^16^. Briefly, for each condition, bacteria were recovered from plates and resuspended in 0.9% NaCl, centrifuged and washed in 0.9% NaCl water solution containing 0.02% Triton X-100, followed by two washes in 0.9% NaCl at 4°C. This washing eliminated lipids from the medium that may adhere to *S. aureus* cells. Cold cell pellets were treated on ice with 0.5 mL of 1N sodium methoxide prepared in methanol. FA methyl esters were extracted after addition of 200 µL of heptane containing methyl-10-undecenoate (Sigma-Aldrich, France) as internal standard. Samples were vortexed for 1 min and centrifuged to separate the phases. FA methyl esters were recovered in the heptane phase (top phase)^16^. Analyses were performed on a Clarus® 580 Gas Chromatograph (Perkin-Elmer) equipped with a Zebron capillary column 30 m x 0.25 mm x 0.25 µm (Phenomenex). Data was analyzed by TotalChrom workstation software (Perkin-Elmer).

### *S. aureus* membrane fluidity measurement

Bacterial membrane fluidity was determined by Laurdan generalized polarization (GP)^31^. An overnight *S. aureus* JE2 and JE2 *ΔfabD* cultures in serum, milk and SerFA were collected, washed once with PBS and incubated in 10 µM Laurdan (Sigma-Aldrich, France) in the dark under agitation at 30°C for 10 min. The stained cells were washed four times with Laurdan buffer (137 mM NaCl, 2.7 mM KCl, 10 mM Na_2_HPO_4_, 1.8 mM KH_2_PO_4_, 0.2 % glucose, 1 % DMF) and transferred to a dark clear-bottom 96-well plates (Corning, France). All solutions and plastics were pre-warmed to 30°C. Laurdan fluorescence was measured between 400 nm and 540 nm under excitation at 350 nm using a Tecan Spark® at 30°C. Laurdan GP was calculated using the formula: GP = (I_440_-I_500_)/(I_440_+I_500_).

### Transmission electron microscopy (TEM)

The JE2 strain was grown to saturation in BHI, serum or milk. Cells were centrifuged (8000 rpm, 5 min) and then resuspended and fixed with 2% glutaraldehyde. Cells were contrasted with 0.5% oolong tea extract in sodium cacodylate buffer as described^56^. Samples were embedded in Epon (Delta Microscopy, Labège, France) and thin sections (70-nm) were collected onto 200-mesh copper grids and counterstained with lead citrate. Grids were observed using a Hitachi HT7700 electron microscope operated at 80 kV (Elexience, France) equipped with a charge-coupled device camera (Advanced Microscopy Techniques Corp., Japan). Bacterial envelope thickness was measured on at least ten cells, taking at least six measurements per cell using ImageJ software. Statistical analysis was performed using Prism 8 (GraphPad Software, San Diego, CA (San Diego, Ca)) with the ANOVA test. A P value of <0.05 was considered significant.

### Imaging of vancomycin binding to *S. aureus*

Imaging of vancomycin binding to *S. aureus* envelopes was performed using BODIPY™ FL-conjugated vancomycin (Invitrogen, France) to *S. aureus* JE2 cells after growth in BHI, serum, or milk. Cells in saturation phase were washed and fixed with 4% paraformaldehyde in PBS, and then incubated with BODIPY™ FL-vancomycin (3 µg/ml) in PBS for 15 min. Subsequently, cells were washed 3 times with PBS and deposited on glass slides. Slides were mounted with Vectashield (Vector Laboratories) and visualized with an Axio-Observer Z1 inverted fluorescence microscope equipped with a Zeiss AxioCam MRm digital camera and Zeiss fluorescence filter, using a Zeiss Apochromat 100x/1.4 oil objective. Images were acquired and processed using the ZEN software package (Zeiss). Quantification of the cells staining was done using open-source imaging software ImageJ.

### Cell permeability assay

*S. aureus* cells grown in serum, milk and BHI overnight were collected and washed with PBS. Pellets were resuspended in PBS and the OD_600nm_ was adjusted to 0.5. A 200 µl aliquot of bacterial suspensions were placed into flat bottom black 96-well plates (Greiner, France), followed by addition of ethidium bromide (EtBr; 0.01 µg/mL final concentration). Once inside, EtBr intercalates into DNA resulting in increased fluorescence emission, and was used essentially as described ^35^. Fluorescence was measured with an excitation and emission wavelength of 530 nm and 585 nm respectively, every 60 seconds for 60 minutes at 37 °C.

### Autolysis assay

To monitor autolysis kinetics, *S. aureus* overnight cultures were diluted to an OD_600_ of 0.025 and grown in different solid media at 37°C. Autolysis assays were conducted on stationary phase cells harvested by centrifugation, washed twice in PBS and resuspended in autolysis buffer (PBS, 0.5% Triton X-100, pH 7.4). The cell suspensions were then incubated at 37°C with shaking and optical densities were recorded every hour during 5h.

### Electrophoretic Mobility and Hydrophobicity

The electrophoretic mobility of JE2 was determined by zeta potential measurements using a Zetasizer Pro (Malvern Panalytical Instrumentation, UK) equipped with a disposable capillary cell (DTS1070). Each sample contained 10^8^ cells/mL, and at least five independent measurements were performed. Results are presented as mean ± SD. The hydrophobicity of the bacterial cell surface was assessed using the microbial adhesion to solvents method, which compares bacterial affinity for a polar solvent (PBS) and a nonpolar solvent (xylene), as previously described by Pinemonti et al. ^35^. Briefly, 300 µL of xylene was added to test tubes containing 2 mL of washed bacterial suspensions (10^8^ CFU/mL) in PBS. The mixtures were vortexed for 120 s and then incubated at 30°C for 10 min to allow phase separation. The aqueous (PBS) phase was carefully collected with a Pasteur pipette and transferred to a UV–Vis cuvette. Optical density was measured at 400 nm using a Novaspect II spectrophotometer (France). All experiments were performed in triplicate using three independent biological replicates.

### Mass spectroscopy: sample preparation, proteolytic digestion, desalting and nanoLC-MS/MS analyses

*S. aureus* JE2 was grown in BHIA, in serum, and in milk plates as above, prepared in 5 independent replicates. Bacteria were collected by centrifugation for 10 min at 7,000 g, resuspended in 200 µL of lysis buffer (8M urea, 25 mM HEPES, pH 8.0, 5% glycerol, 1mM DTT, 0.2% DDM (n-dodecyl-β-D-maltopyranoside), 1:200 v/v protease inhibitor (Thermo Fisher Scientific, Canada) and homogenized (using vortex mixer) for 5 min. The suspension was centrifuged for 10 min at 15,000 g and the soluble protein extract was separated from the insoluble debris. Protein concentrations were measured by the Bradford assay using a protein assay kit (Thermo Fisher Scientific) following the manufacturer’s protocol. The assay absorbance was measured with a spectrophotometer (Novaspec III, Biochrom) and semi-microvolume disposable polystyrene cuvettes (Bio-Rad) at 595 nm.

Protein extracts (100 µg) were digested using a modified filter-aided sample preparation (FASP) protocol designed for proteomic analysis^71^. The digested samples were desalted on disposable Pierce C-18 tips (Thermo Fisher Scientific) with the addition of C-18 resins from Pierce C-18 Spin Columns (Thermo Fisher Scientific). Samples were then dried by vacuum centrifugation (Savant SPD111V SpeedVac Concentrator, Thermo Scientific) and reconstituted with MS grade water with 0.1% formic acid for MS analysis.

NanoLC-MS/MS analyses were performed as described with some modifications of liquid chromatographic gradient on an Ultimate3000 nanoRLSC (Thermo Scientific) coupled to an Orbitrap FusionTM (Thermo Scientific)^72^. Two µL of protein digests were injected and separated on a column (15 cm LT x 75 μm i.d. x 365 μm o.d. fused silica capillary, Polymicro Technologies) packed in-house with Luna C18 particles (Luna C18(2), 3 μm, 100 Å, Phenomenex, Torrance, California, USA). The mobile phase consisted of a mixture of water/ACN/0.1% (v/v) FA, working at a flow rate of 0.30 μL/min (0-7 min, 2-2% ACN; 7-107 min, 2-38% ACN; 107-112 min, 38-98% ACN; 112-122 min, 98-98% ACN; 122-130 min, 98-2% ACN; 130-140 min, 2-2% ACN).

### Data analysis

MaxQuant (Thermo Scientific, version v1.6.17.0)^73,74^ was used for protein identification and quantification. A maximum of two missed tryptic cleavages were allowed with precursor charge bounded between +2 and +7, a 10ppm precursor mass tolerance, and a 0.5 Da fragment mass tolerance. Moreover, search parameters were set to allow for dynamic modifications, including methionine oxidation, acetylation on the N-terminus, and fixed cysteine carbamidomethylation. The search database consisted of a non-redundant protein sequence FASTA file containing the 3,936 entries from *S. aureus* strain USA300 - Uniprot taxonomy ID 367830 (2024.10.09). Normalized label-free quantification (LFQ) values were obtained by applying the in-built MaxLFQ algorithm^73^.

Bioinformatics of sample LFQ intensities was conducted using R programming language in RStudio. Potential contaminants were removed and proteins that were found in at least 4/5 biological replicates were used for further analysis. To visualize common and unique proteins, a Venn diagram was created using the VennDiagram package in R where unique proteins were defined as proteins that were present in 4/5 samples of a particular group and absent in at least 3/5 samples of all other groups. Additionally, proteins common to all three groups were defined as those present in 4/5 samples of each group. To comparatively visualize the whole proteome, a principal component analysis was conducted. Missing LFQ intensity data was first imputed. The data was then log2 normalized and scaled. The components were then calculated using the prcomp() function from the stats package. Principal component 1(PC1) and principal component 2(PC2) were plotted using the ggplot2 package. The data used to make the PCA plot was then used to make the heatmap with the heatmap package in R.

Additionally, a GroupWise comparison of the log_2_ normalized data was performed of common proteins by conducting a one-way Anova with the aov() function from the stats package. A Tukey’s Honest Significance Difference (TukeyHSD) test was then used to adjust the p-values. In addition to the adjusted p-values, the log2 fold change was also calculated. Both parameters, comparing serum *versus* BHI, milk *versus* BHI, and serum *versus* milk, were plotted to create volcano plots. Overall protein significance was defined as having an adjusted p value of <0.05 and a minimum FC of 1.5.

### Staphyloxanthin pigment quantitation by spectroscopy

Spectral profiles of *S. aureus* pigment were assessed as described^50^. Bacterial cells of the JE2 strain were recovered after 48h growth at 37°C on 20% milk, 20% serum, and BHI agar plates, and adjusted to OD_600_ = 2 for all samples. For methanol extractions, bacteria were washed with milliQ water, pelleted and resuspended with 99% methanol. Cell suspensions were incubated for 5 min in a 55°C water bath, and then cooled 10 min at room temperature. Supernatants were collected after centrifugation 15 min at 6,000 rpm. Absorbance profiles of the extracted carotenoids were determined using the Cary 3500 Multicell UV-Vis Spectrophotometer (Agilent, USA).

### ROS sensitivity assays

Tests for *S. aureus* susceptibility to oxidants were performed in PBS as described^50^. For the singlet oxygen assay, 10^8^ *S. aureus* cells were incubated at room temperature in individual wells of a 24-well culture plate in the presence of 6 μg/ml methylene blue. The plate was situated 5 cm from a 100 W visible light source. Bacterial viability was assessed after 3 hr by plating dilutions on BHIA. Control plates were handled identically but exposed to light in the absence of methylene blue; no bacterial killing was observed in controls. For hydrogen peroxide (H_2_O_2_) tests, H_2_O_2_ (1.5 % final concentration) was added to 2 × 10^9^ CFU *S. aureus* suspensions. H_2_O_2_ was replaced by water in controls. Samples were incubated at room temperature for 1 hr, and then 1,000 U/ml of catalase (Sigma-Aldrich, France) was added to quench residual H_2_O_2_. Milk-adapted and serum-adapted cells were washed and resuspended in PBS prior to ROS challenge to assure identical test conditions. Dilutions were plated on BHIA for enumeration of surviving cells.

### Antibiotic resistance

Minimal inhibitory concentrations (MIC) of nisin, ciprofloxacin and vancomycin were determined by microdilution tests for *S. aureus* JE2 and *crtM* strains. Overnight cultures in serum, milk and BHI agar plates were collected and washed in PBS, and bacterial OD_600_ was adjusted to 0.5 in the same buffer. Two-fold serial antibiotic dilutions were generated in U-bottom 96-well plates (Greiner, France). Wells were inoculated with bacterial suspensions and incubated 16h at 37 °C with 180 rpm shaking. Bacterial viability was assessed by plating dilutions on BHI agar. The MIC was defined as the lowest concentration of antibiotic that inhibited growth.

### *G. mellonella* virulence assay

*G. mellonella* larvae (weighing ∼250 mg) were reared in INRAE facilities (Jouy en Josas, France) on pollen and bee wax (La Ruche Roannaise Besachier, France). The rearing container was maintained at 27°C in a humidified incubator. A total of 200 last-instar larvae used for the experiment were subjected to starvation for 24 hr at 27°C prior to infection. Based on previous CFU determinations, 10^6^ CFU *S. aureus* per larva was determined as an optimal inoculum required to kill *G. mellonella* larvae within 72 hr ^9^. An overnight *S. aureus* preculture in BHI, serum and milk were subcultured on corresponding solid plates by spreading a 100 µl aliquot for overnight growth. Each culture was collected, washed 3 times in PBS, and then resuspended in PBS to obtain 10^8^ CFU/ml. Larvae were infected by intrahemocoelic injections with 10 μl of bacterial suspensions using a syringe and needle with an injector pump KDS100 (KD Scientific, ThermoFisher, Illkirch, France). Inocula were counted after plating onto BHI agar medium. Control larvae were injected with saline alone. Larvae were incubated in 9 cm diameter plastic dishes (10 per dish) without food at 37 °C. Mortality was scored at 24-, 48-, and 72-hour post-infection. At least 20 larvae were infected per condition. Results are presented as the mean ± standard deviation (SD).

### Statistics

The significance of experimental differences in oxidant sensitivity, envelope thickness, cell size and membrane fluidity were evaluated by ANOVA test. Results of the insect in vivo challenge studies were evaluated by nonparametric t-test (Mann-Whitney U). The data were analyzed using Prism V8 software (GraphPad). CLSM images were analyzed by ImageJ (v2.9.0; 64-bit).

## Contributions

A.G., B.H., and J.V. designed the study, provided materials and funding. V.L., K.G., Z.M., R.D’M., C.N.L., A.G., and J.V.: performed experiments, analyzed the data, and participated in writing the manuscript. M.K., M.C., C.P., M.S., S.T.: performed experiments, analyzed the data. V.L., Z.M., C.N.L., M.M., K.G., P.G., A.G. and J.V. wrote and revised the manuscript. All authors discussed the manuscript.

## Data availability

The mass spectrometry proteomics data have been deposited to the ProteomeXchange Consortium via the PRIDE partner repository with the dataset identifier PXD064173.

## Acknowledgments

We are grateful to Jamila Anba-Mondoloni, Francesco Rizzotto, and Clara Louche (INRAE, France) for stimulating discussion, and Yingxi (Cici) Li (University of Ottawa, Canada) and Christophe Buisson (INRAE, France) for valuable technical assistance. We acknowledge the INRAE MIMA2 imaging platform https://doi.org/10.15454/1.5572348210007727E12 and the University Ottawa bioinformatics facility.

## Funding

This work was funded by the Agence Nationale de Recherche (ANR-21-CE21-009, Siena and ANR-21-CE42-008, Elise to JV, ANR-16-CE15-0013 to AG and under the umbrella of the Joint Programming Initiative on Antimicrobial Resistance (JPIAMR) ANR-22-AAMR-0007 to AG), Fondation pour la Recherche Médicale (DBF20161136769 to AG), Ministry of Science, Technological Development and Innovation of the Republic of Serbia (451-03-136/2025-03/200026 to MS) and in part by the European Union under grant agreement N° 101135402 (MOBILES project to JV), and under grant agreement N° 872662 (IPANEMA project to JV). Vincent Léguillier was recipient of a PhD grant funding from the Région Ile-de-France (DIM 1Health2021).

## Notes

https://www.ebi.ac.uk/pride/archive/projects/PXD064173/

### Competing Interest Statement

The authors have declared no competing interest.

