## Supplementary Information for "Biotope-dependent High Level Resistance to Reactive Oxygen Species, Antibiotic Tolerance, and Virulence of *Staphylococcus aureus*"

Supporting Information description: additional experimental details

**Table of content Summary**

**Supplementary Figure 1-** *S. aureus* Δ*fabD* mutant is a fatty acid (FA) auxotroph.
**Supplementary Figure 2-** *S. aureus* Δ*fabD* mutant growth in 100% adult bovine serum and 100% raw cow milk.
**Supplementary Figure 3-** Cross-sections of *S. aureus* JE2 cultures in BHI, serum, and milk medium examined by Transmission Electron Microscopy (TEM).
**Supplementary Figure 4-** BODIPY™ FL- vancomycin staining of *S. aureus* JE2 cultures in BHI, serum, and milk.
**Supplementary Figure 5-** Overview of proteomics results.
**Supplementary Figure 6-** Global changes of proteins extracted from saturated *S. aureus* cultures.
**Supplementary Figure 7-** Volcano plots of quantified proteins in serum- and milk- *versus* BHI-grown *S. aureus* JE2.
**Supplementary Figure 8-** Proteomic reprograming in milk- and serum-adapted *S. aureus*.
**Supplementary Figure 9-** Singlet oxygen stress on serum- and milk-adapted *S. aureus* RN4220 derivative RN-R.
**Supplementary Figure 10-** Hydrogen peroxide stress on serum- and milk-adapted *S. aureus* JE2.
**Supplementary Figure 11-** Sensitivity of *S. aureus* against ciprofloxacin, vancomycin and nisin in the presence of ascorbic acid.

**Supplementary Table 1 -** Fatty acid composition of JE2 and Δ*fabD* grown in adult bovine serum or milk.
**Supplementary Table 2-** Bacterial strains used in the study.
**Supplementary Table 3 -** Primers used for Δ*fabD* mutant construction.

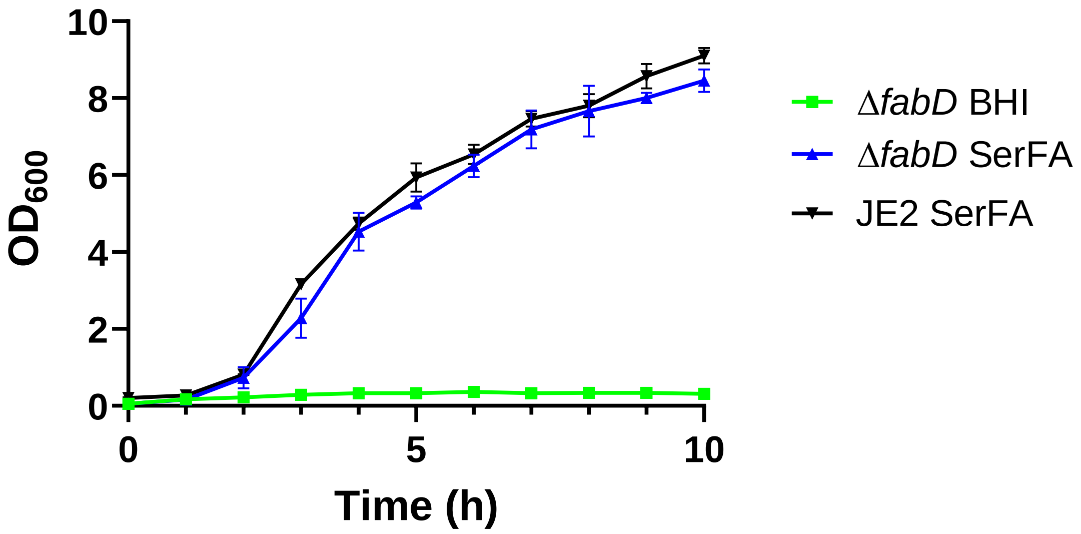


**Supplementary Fig. 1. The *S. aureus* Δ*fabD* mutant is a fatty acid (FA) auxotroph.** FabD (malonyl CoA-acyl carrier protein transacylase) initiates the FA synthesis pathway II (FASII), and *fabD* mutants are fatty acid auxotrophs [1,2]. The *S. aureus* Δ*fabD* mutant cannot multiply in BHI, but grows nearly as well as the parental JE2 strain when grown in SerFA (BHI containing 10% serum and 0.17 mM each of myristic acid (C14:0), palmitic acid (C16:0) and oleic acid (C18:1); note that serum neutralizes FA toxicity[3].

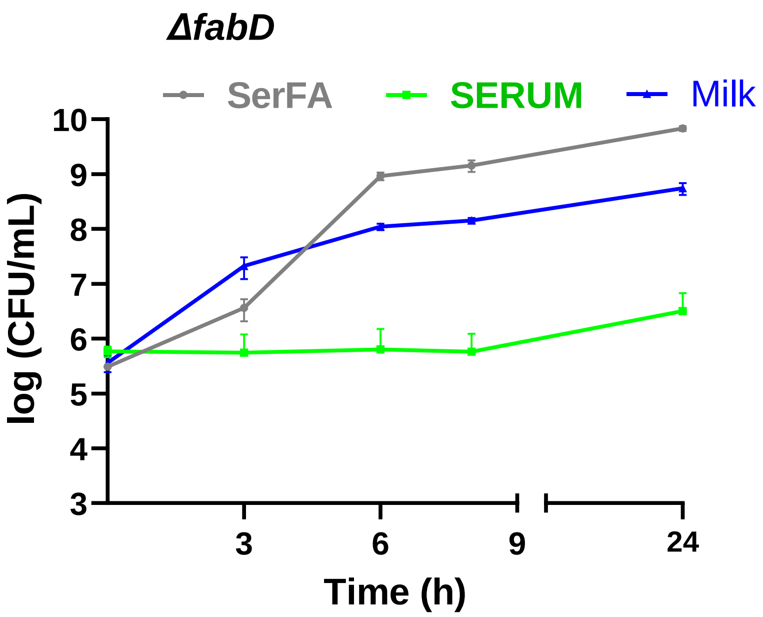


**Supplementary Fig. 2. *S. aureus* Δ*fabD* mutant growth in 100% adult bovine serum and 100% raw cow milk.** Growth of *ΔfabD* mutant in serum and milk is compared to its growth in SerFA medium. Data presents mean ± SD of three biological replicates.

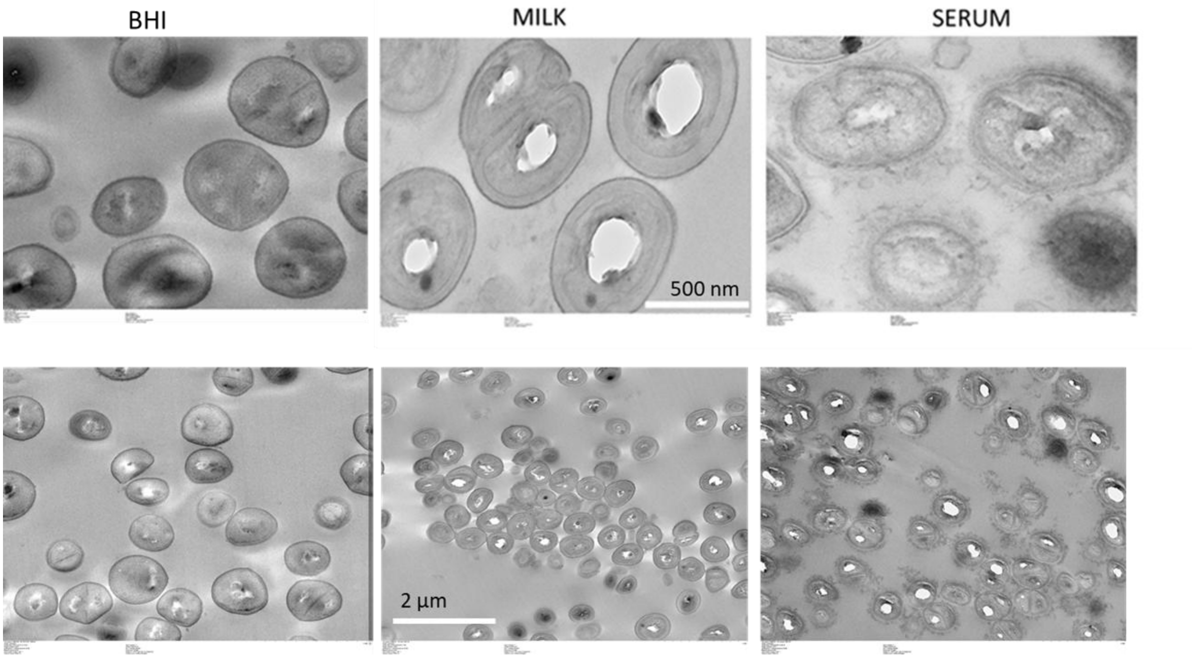


**Supplementary Fig. 3. Cross-sections of *S. aureus* JE2 cultures in BHI, serum, and milk medium examined by Transmission Electron Microscopy (TEM).** Scale bars are at 500 nm (upper images) and 2 µm (lower images).

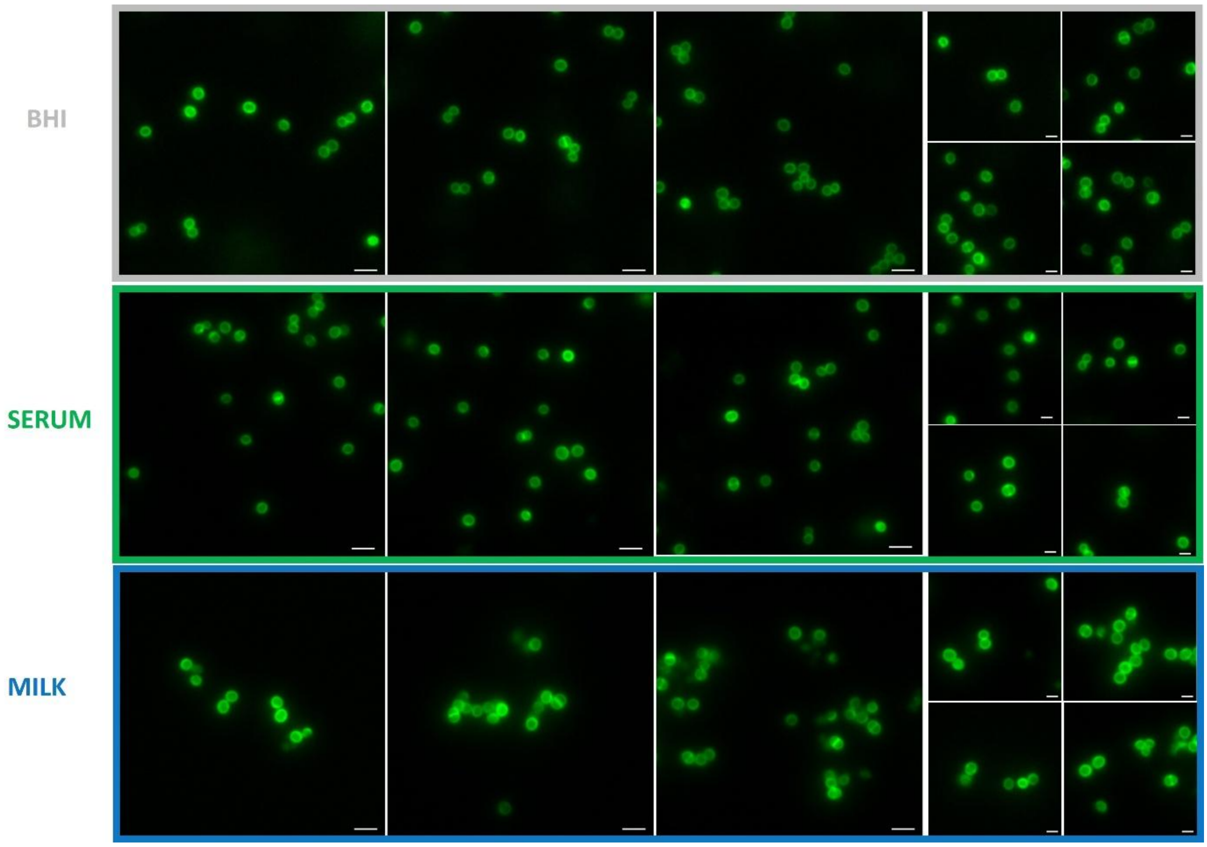


**Supplementary Fig. 4. BODIPY™ FL- vancomycin staining of *S. aureus* JE2 cultures in BHI, serum, and milk medium.** Bacterial cultures were grown in BHI (control cells) or adapted to serum, and milk. Stationary phase bacterial cultures in BHI, serum, and milk, were labeled with BODIPY™ FL- vancomycin and fluorescence at 488 nm was monitored using a fluorescence.

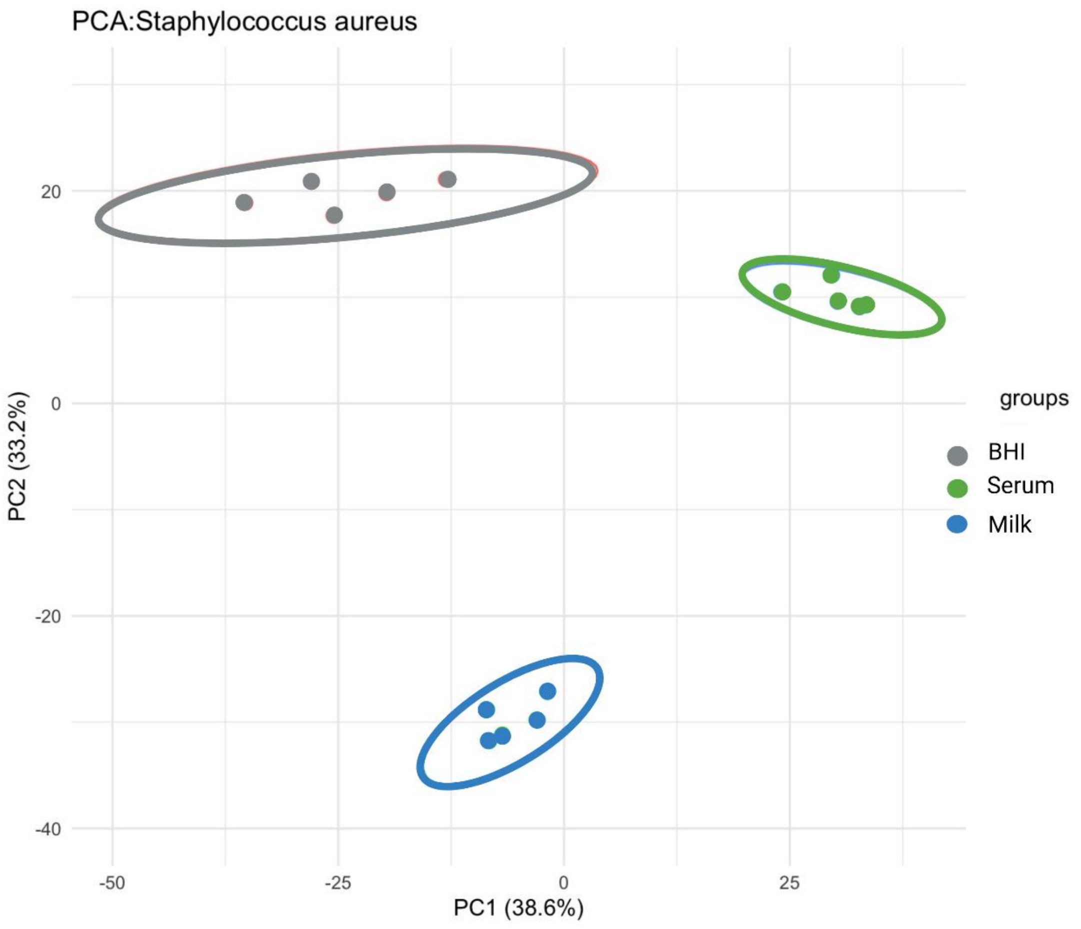


**Supplementary Fig. 5. Overview of proteomics results.** Principal component analysis (PCA) plot depicting the proteomics results of saturated *S. aureus* JE2 cultures in BHI (red), serum (blue), and milk (green) medium. Each dot represents one sample and ellipses indicate 95% confidence areas for each bacterial sample.

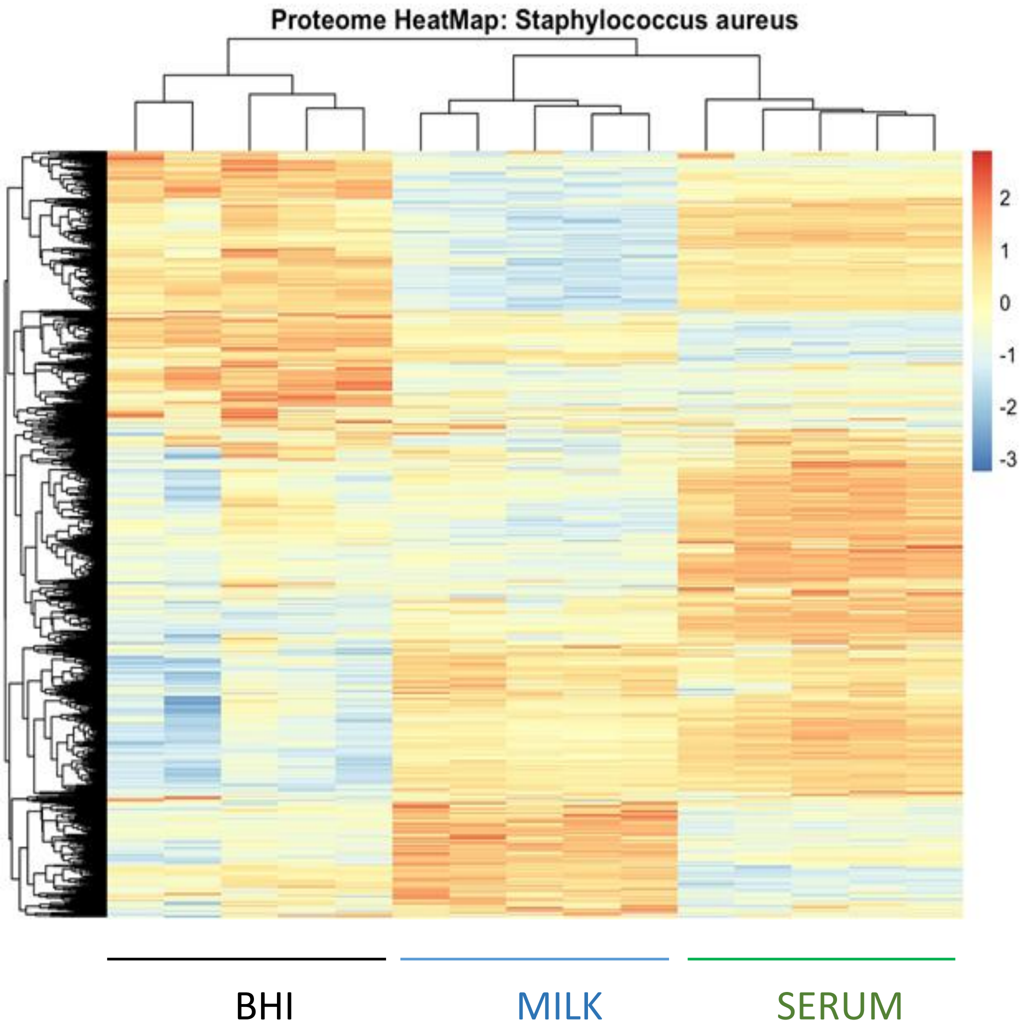


**Supplementary Fig.6. Global changes of proteins extracted from saturated BHI, serum, and milk *S. aureus* JE2 cultures.** Clustering of protein abundance across experimental conditions. Each column presents predominance of the identified proteins and their distribution in each sample. Data shows results of 5 independent replicates for each condition.

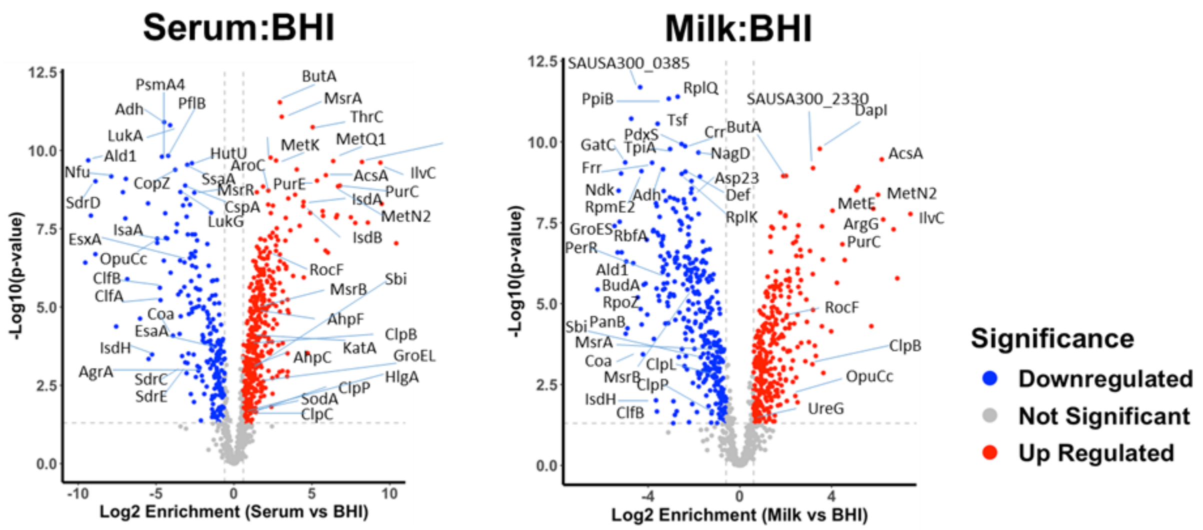


**Supplementary Fig. 7. Volcano plots of quantified proteins extracted from serum- and milk - *versus* BHI- grown *S. aureus* JE2.** Red dots indicate significantly increased proteins, and blue spots indicate significantly downregulated proteins, when compared to levels in BHI conditions; grey dots represent proteins that did not change significantly between the compared groups; labelled proteins show significant differences between conditions. The analysis threshold was p-value < 0.05 and the fold change (FC) > 1.5 or < 0.67; values below these thresholds were not retained as significant. Results of the full proteome analyses are available in **Supplementary Data 1**.

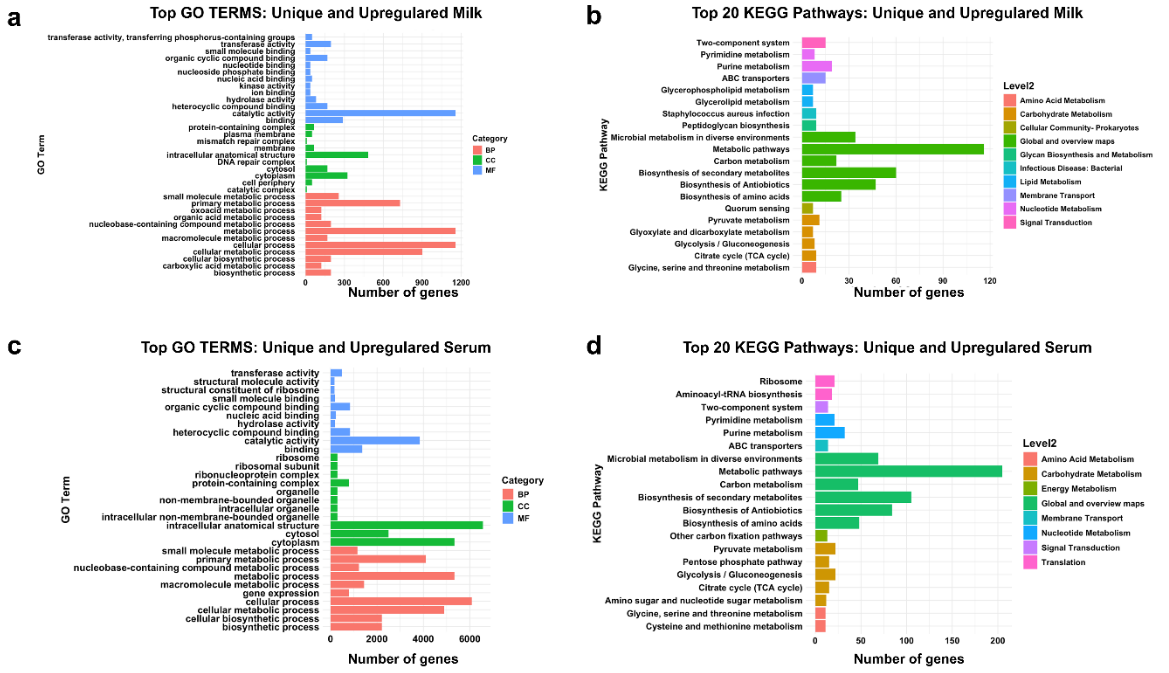


**Supplementary Fig. 8. Proteomic reprogramming in milk- and serum-adapted *S. aureus*.** Gene Ontology (GO) term enrichment analysis of detected differentially expressed proteins ($P < 0.05$) for molecular function (MF), biological process (BP) and localization of detected proteins in cell compartments (CC) between milk-adapted JE2 cells vs. cells cultured in BHI (a) and serum- *versus* BHI-cultured bacteria (b). Bar plot graphs for Kyoto Encyclopedia of Genes and Genomes (KEGG) functional annotation of the differently expressed proteins in milk-adapted JE2 compared to controls in BHI (c) and serum-adapted compared to BHI (d) with a $P_{adj}$ threshold cut-off of 0.05. The longer bar indicates greater protein enrichment; colors indicate the Level 2 subcategories within the pathway-related processes or functions.

Thirty-one stress-response proteins and 49 virulence-related proteins were differentially expressed according to conditions (Supplementary Data 2). The stress proteins participate mainly in adaptation to pH. Numerous virulence-related proteins were up-regulated in milk, and included ≥ 2-fold greater levels of Type VII secretion system and ESAT-6 secretion machinery proteins compared to both serum-adapted and BHI grown cells (Fig. 4e). In contrast, levels of numerous surface proteins Spa, ClfA/B, SdrD/E, Coa were overall lower in both serum- and milk-adapted cells compared to those grown in BHI. Their reduced expression could alter *S. aureus* surface charge and polarity in those conditions (Figs. 3d and 3e), potentially influencing their adhesion capabilities and overall pathogenicity.

Differentially expressed proteins are organized according to corresponding GO biological process (BP), cell component (CC) and molecular function (MF) categories. Remarkably, the top unique and upregulated proteins in serum-adapted cells involved approximately six times more proteins than those inventoried in milk-adapted cells, and were in the CC category of intracellular anatomic structure, cytosol and cytoplasm, including stress-response functions. This observation suggests that serum is a more stressful environment for *S. aureus* than milk, which was experimentally confirmed (see Fig. 1a).

KEGG analysis revealed that metabolic and biosynthetic pathways were highly enriched in both milk- and serum-adapted JE2 compared to BHI (c, d). Enrichment of ABC transporters and 2-component systems indicates adjustments of in transmembrane traffic of substances needed for bacteria growth in milk and serum[4]. Milk-adapted cells showed specific enrichment in infection, glycerolipid metabolism, peptidoglycan biosynthesis, and quorum sensing pathways, which may suggest enhanced virulence. In contrast, proteins involved in aminoacyl-tRNA biosynthesis and pentose phosphate pathways were enriched in serum-adapted but not in milk-adapted cells. These pathways are involved in adjustments to altered environmental conditions[5], and NADPH production, respectively, and may be up-regulated in hostile environments[6]. Overall, these condition-specific responses suggest that milk provides a non-stressful environment for *S. aureus*, whereas serum imposes environmental stress such that survival is prioritized.

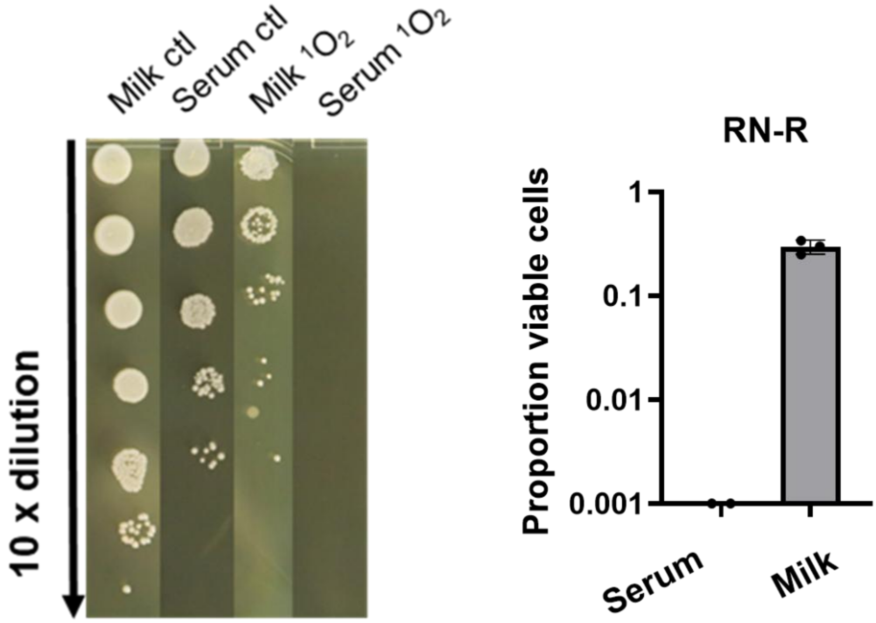


**Supplementary Fig. 9. Singlet oxygen stress on serum- and milk-adapted *S. aureus* RN4220 derivative RN-R.** After exposure to singlet oxygen, cultures of RN-R, an RN4220 strain whose *fakB1* gene was repaired[7] were serially diluted, spotted, and grown overnight to estimate survival (left panel shows representative results of three experiments. Lanes shown correspond to experiments done in parallel but plated on different plates. The mean ± SD based on 3 experiments are shown (right panel). Note that single oxygen treatment eradicated serum-adapted RN-R, but only partially killed the milk-adapted cells.

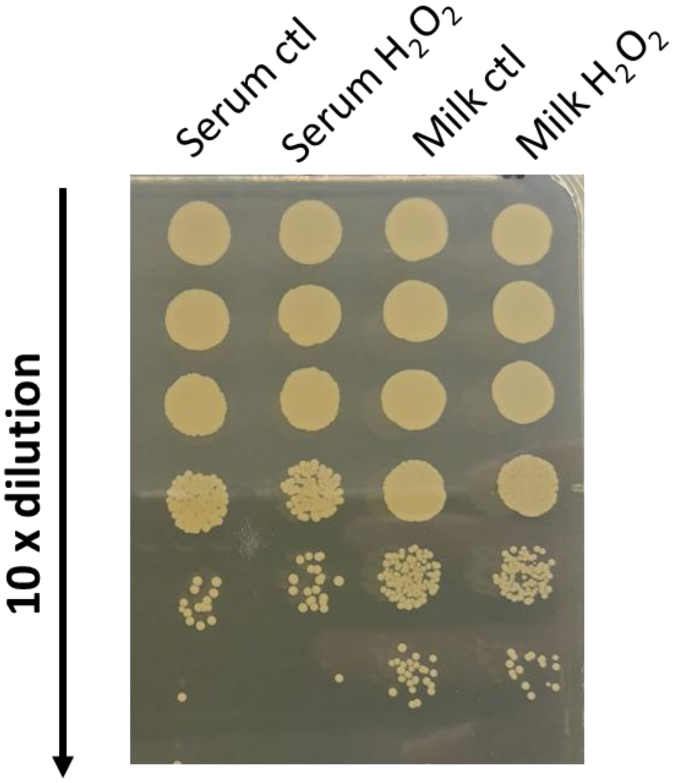


**Supplementary Fig. 10. Hydrogen peroxide stress on serum- and milk-adapted *S. aureus* JE2.** After exposure to 1.5% $H_2O_2$ in PBS, cultures of serum- and milk-adapted JE were serially diluted, spotted on the BHI plate, and grown overnight to estimate survival. Note that no reduction in cell survival compared to the corresponding control was observed. Results is representative of 3 independent experiments.

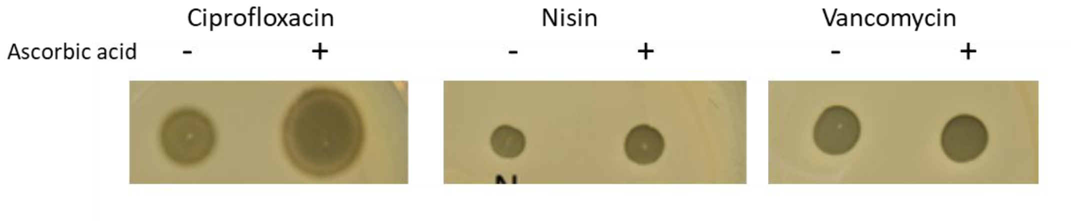


**Supplementary Fig. 11. Sensitivity of *S. aureus* against ciprofloxacin, vancomycin and nisin in the presence of ascorbic acid**. Overnight BHI culture of JE2 was washed in BHI and plated on BHI solid medium. Plates were spotted with 1.25 ng nisin (N), 400 ng ciprofloxacin (C), 10 µg vancomycin (V), 1.25 ng nisin+10 mM ascorbic acid (NC), 400 ng ciprofloxacin + 10 mM ascorbic acid (CC), and 10 µg vancomycin + 10 mM ascorbic acid (VC), respectively. The result shown is representative of three biological replicates.

**Supplementary Table 1. Fatty acid composition of JE2 and Δ*fabD* grown in adult bovine serum or milk.**

| | | | | | | | |
|---|---|---|---|---|---|---|---|
| | | SERUM | | | MILK | | |
| | | Serum | JE2 - Serum | Δ*fabD* - Serum | Milk | JE2 - Milk | Δ*fabD* - Milk |
| Saturated FA | 14:0 | 0.5 | 1.0 | 0.0 | 14.3 | 11.7 | 25.8 |
| | ai15 | 0.0 | 6.6 | 0.0 | 0.6 | 15.5 | 9.9 |
| | 16:0 | 16.0 | 11.9 | 19.2 | 48.0 | 13.2 | 19.4 |
| | 17:0 | 5.7 | 3.7 | 4.8 | 0.0 | 0.0 | 0.0 |
| | 18:0 | 24.7 | 17.1 | 18.4 | 13.9 | 18.7 | 0.8 |
| | 20:0 | 0.0 | 2.9 | 0.0 | 0.0 | 7.6 | 0.0 |
| Unsaturated FA | 18:1 | 18.3 | 22.2 | 20.9 | 22.6 | 22.5 | 44.0 |
| | 18:2 | 22.0 | 20.8 | 22.8 | 0.0 | 0.0 | 0.0 |
| | 18:3 | 8.0 | 9.0 | 9.2 | 0.7 | 0.0 | 0.0 |
| | 20:1 | 0.0 | 1.5 | 0.0 | 0.0 | 10.7 | 0.0 |
| | 20:3 | 2.3 | 1.5 | 2.5 | 0.0 | 0.0 | 0.0 |
| | 20:4 | 2.6 | 1.9 | 2.1 | 0.0 | 0.0 | 0.0 |
| | | 100.0 | 100.0 | 100.0 | 100.0 | 100.0 | 100.0 |
| | Saturated | 46.8 | 43.1 | 42.5 | 76.7 | 66.8 | 56.0 |
| | Unsaturated | 53.2 | 56.9 | 57.5 | 23.3 | 33.2 | 44.0 |

**Supplementary Table 2. Bacterial strains.**

| Strain | Description | Reference or Source |
|---|---|---|
| JE2 | FPR3757 USA300-JE2 (referred to as JE2) | [8] |
| Δ*fabD* | US300 with the *fabD* gene deleted | This study |
| RN-R | RN4220 *S. aureus* cloning recipient ATCC 8325-4 derivative restriction negative | [9] |
| JE2 *katA* mutant | USA300 derivative lacking the katalase A | BEI Resources, [8] |
| JE2 *crtM* mutant (*crtM*::Tn, SAUSA300_2499) | USA3000 derivative lacking the fatty acid kinase A | BEI Resources, [8] |
| *E. coli* DH5α | Used for Δ*fabD* construction | [10] |

**Supplementary Table 3. Primers used for *ΔfabD* mutant construction**

| Primer Name | Primer sequence (5' – 3') |
|---|---|
| pMad_F | CGAATTCTAGAAGCTTCTGC |
| pMad_R | CTGTCTAGTTAATGTGTAACG |
| PlsX_F | CTGAACACTTATTACAGTATGC |
| FabG_R | AACTGCATTTACAGTGATACC |
| pMad-plsX | cgatgcatgccatggtacccATTAGAGGTATTGATAATCCG |
| plsX_FabG_R | CCTGTTACTAAAGCACTCTTAGTCATTTTACTCATTTGATTCACCTACAG |
| plsX-FabG_F | CTGTAGGTGAATCAAATGAGTAAAATGACTAAGAGTGCTTTAGTAACAGG |
| fabG_pMad_R | gcttctagaattcgagctcccGCCGCAGATTTAGTTAAACC |

1 Gloux, K. *et al.* Clinical relevance of type II fatty acid synthesis bypass in Staphylococcus aureus. *Antimicrobial agents and chemotherapy* **61**, e02515-02516 (2017).
2 Morvan, C. *et al.* Environmental fatty acids enable emergence of infectious Staphylococcus aureus resistant to FASII-targeted antimicrobials. *Nature communications* **7**, 1-11 (2016).
3 Kénanian, G. *et al.* Permissive fatty acid incorporation promotes staphylococcal adaptation to FASII antibiotics in host environments. *Cell reports* **29**, 3974-3982. e3974 (2019).
4 Bronner, S., Monteil, H. & Prévost, G. Regulation of virulence determinants in Staphylococcus aureus: complexity and applications. *FEMS microbiology reviews* **28**, 183-200 (2004).
5 Pelz, A. *et al.* Structure and biosynthesis of staphyloxanthin from Staphylococcus aureus. *Journal of Biological Chemistry* **280**, 32493-32498 (2005).
6 Liu, Y. *et al.* A bacterial pigment provides cross-species protection from H2O2-and neutrophil-mediated killing. *Proceedings of the National Academy of Sciences* **121**, e2312334121 (2024).
7 Pathania, A. *et al.* (p) ppGpp/GTP and malonyl-CoA modulate Staphylococcus aureus adaptation to FASII antibiotics and provide a basis for synergistic bi-therapy. *Mbio* **12**, e03193-03120 (2021).
8 Fey, P. D. *et al.* A genetic resource for rapid and comprehensive phenotype screening of nonessential Staphylococcus aureus genes. *MBio* **4**, e00537-00512 (2013).
9 Kreiswirth, B. N. *et al.* The toxic shock syndrome exotoxin structural gene is not detectably transmitted by a prophage. *Nature* **305**, 709-712 (1983).
10 Sambrook, J., Fritsch, E. F. & Maniatis, T. *Molecular cloning: a laboratory manual*. (Cold spring harbor laboratory press, 1989).
